# Genomic insights into host shifts between *Plasmodium vivax* and *Plasmodium simium* in Latin America

**DOI:** 10.1101/2024.12.19.629455

**Authors:** Margaux J. M. Lefebvre, Fanny Degrugillier, Céline Arnathau, Camila González, Silvia Rondón, Andrés Link, Andrea Chaves, Julio A. Benavides, Aline Alves Scarpellini Campos, Edmilson dos Santos, Rosana Huff, Cláudia Maria Dornelles Silva, Ezequiel Vanderhoeven, Benoit De Thoisy, Michael C. Fontaine, Franck Prugnolle, Virginie Rougeron

## Abstract

Malaria in Latin America is largely caused by *Plasmodium vivax*, but its lesser-known sister species, *Plasmodium simium*, has recently emerged from monkeys to infect humans, thus raising new public health concerns. By analyzing 719 monkey samples and whole genome variations for 19 *P. simium* and 408 *P. vivax* isolates, we investigated the evolutionary history and population genetics of the two species. *P. vivax*, typically restricted to humans, was identified in three Colombian and one Brazilian monkeys, suggesting host niche expansion. Genetic analysis reveals recent genetic exchanges between both species and indicates that *P. simium* originated from a host jump approximately a century ago, possibly linked to *P. vivax* migration from Mexico to Brazil. Genome-wide scans revealed signals of positive selection in *P. simium* genes involved in interactions with primate hosts and mosquito vectors. These findings highlight *P. simium* evolutionary history and zoonotic malaria risks, and underscore the need to include monkeys in malaria prevention measures while ensuring human-wildlife coexistence.

## Introduction

*Plasmodium* species are responsible for malaria in a wide range of vertebrate hosts, including primates, rodents, reptiles and birds (*1*). Within this genus, *Plasmodium falciparum* and *Plasmodium vivax* are the most clinically significant species in terms of malaria-related morbidity and mortality in humans (*2*). Other species, such as *Plasmodium malariae*, *Plasmodium ovale curtisi*, *Plasmodium ovale wallikeri*, *Plasmodium knowlesi*, *Plasmodium cynomolgi* and *Plasmodium simium*, also can infect humans, contributing to the complexity of malaria epidemiology (*3*). *P. knowlesi*, *P. cynomolgi*, and *P. simium* primarily infect non-human primates (NHPs), but also humans, contributing to emerging zoonotic malaria cases (*4–6*). For example, *P. knowlesi* and *P. cynomolgi*, which naturally infect Asian NHPs, have emerged as a significant public health issue in South-East Asia (*7–9*). *P. simium* has more recently been implicated in human malaria cases in Latin America (Brazil) (*5*). Today, zoonotic malaria, caused by malaria parasites transmitted from animals to humans, is becoming problematic due to human encroachment on forest habitats (*10*, *11*). The increased risk of transmission from NHP reservoirs (*10*, *11*) potentially challenges malaria elimination programs and makes zoonotic malaria a potential public health issue.

In South America, *P. simium* primarily infects platyrrhine monkeys (*12*). It was initially described in 1951 from blood smears of a southern brown howler monkey (*Alouatta clamitans*) near São Paulo, Brazil (*13*), and since then it has been detected in other monkey species, including the howler monkeys (*Alouatta fusca* and *Alouatta caraya*), southern muriqui (*Brachyteles arachnoides*), three-striped night monkeys (*Aotus trivirgatus*), and capuchin monkeys (*3*, *14*). Its zoonotic potential was first described in 1966 when human infections were identified in Brazil through blood smear analysis, raising concerns about cross-species transmission (*15*). Historically, *P. simium* is believed to be endemic in Brazil (*5*, *12*, *16*). The detection of infections in NHPs from Colombia in 2019 (*17*) and Costa Rica in 2022 (*18*) suggested a broader distribution, although it remains unclear whether these infections were caused by *P. simium* or *P. vivax*.

Genetically, *P. simium* is closely related to *P. vivax*, although they can be distinguished by two mitochondrial single nucleotide polymorphisms (SNPs) and morphological differences in erythrocytic stages (*5*, *19*). In addition, no hybrid has been identified to date (*20*), supporting the classification of *P. simium* as a distinct species from *P. vivax* (*13*). However, due to their genetic similarities, shared host range (humans), and potential for recombination, the distinction between *P. simium* and *P. vivax* may not be as clear-cut as currently understood and further investigation are required to clarify the taxonomic status of the two species.

*Plasmodium simium* evolutionary origin(s) and history was largely overlooked (*3*) until 2017, when Brasil *et al.* (*5*) characterized its mitochondrial genome following a human malaria epidemic in Brazil. They showed that this epidemic was caused by *P. simium* isolates. Then whole genome sequencing (WGS) studies confirmed that *P. simium* is genetically close to the American human *P. vivax* and characterized by a lower genetic diversity (*20*, *21*), suggesting a host jump to Neotropical NHPs (*14*, *20*, *21*) (*i.e.,* reverse zoonosis (*21*)). Intriguingly, some studies showed that Brazilian *P. simium* strains are more closely related to Mexican than to Brazilian *P. vivax*, without providing any explanation for such results (*20*, *21*). Other studies, based on mitochondrial genomes and some nuclear markers, suggested the occurrence of multiple host jumps (*22*, *23*), that involved not only human *P. vivax* populations from Latin America, but also potentially from Asia (*24*, *25*). Additionally, although the genetic proximity of these species suggests a recent host jump and thus a recent divergence (*12*, *14*, *20*, *21*), to our knowledge, no precise date for this event has been provided. *P. simium* evolutionary history and origin(s) remain thus still unresolved.

The mechanisms by which *P. simium* has adapted to different hosts and vectors are also poorly understood. No study has explored the population genomics of the host jump from humans to NHPs. This is likely to have exerted strong selective pressures on the parasite, driving its adaptation to new hosts and resulting in distinct phenotypes and genotypes. Only some studies targeting a few candidate genes identified some genes that may have facilitated *P. simium* successful invasion of the Neotropical primate hosts (*20*, *21*). For instance, deletions were found in the coding regions of *pvrbpa2* (reticulocyte binding protein 2a) and *pvdbp1* (Duffy binding protein 1), two genes involved in the invasion of reticulocytes in primates (*20*, *21*). Other than adapting to primate hosts, the parasite has also to adapt to the vector responsible for its transmission. In Latin America, *P. vivax* is mainly transmitted by *Anopheles darlingi* (*26*, *27*) and *Anopheles albimanus* (*27*, *28*). Conversely, it is thought that *P. simium* is transmitted by anopheline mosquitoes of the *Kerteszia* subgenus, primarily *Anopheles cruzii* and *Anopheles bellator* (*29*, *30*), suggesting adaptations to these distinct mosquito vectors. For example, *Pvs47*, a gene involved in the evasion of the mosquito immune system, harbors two amino acid changes in *P. simium*, but absent in global *P. vivax* populations (*21*). However, a comprehensive genomic understanding of these adaptations is still lacking.

Given the zoonotic potential of *P. simium*, understanding its geographic range, evolutionary origin(s), and adaptation mechanisms is fundamental for developing effective malaria control and elimination strategies, but also for the conservation of endangered NHPs that might be affected by this pathogen. The aim of this study was to clarify *P. simium* geographic distribution and host range in Latin America and to investigate its evolutionary history and adaptations. To address these questions, we first screened 719 NHP samples from five Latin American countries (Colombia, Brazil, Argentina, Costa Rica and French Guiana) for the presence of *P. simium*. This extensive screening identified three Colombian NHPs infected with *P. vivax* and not *P. simium*, a finding that, to our knowledge, has not been previously reported. We then analyzed whole genome variations for these samples and previously published *P. simium* (n=19) and *P. vivax* (n=405) datasets to assess population genetic diversity, structure, evolution history, and evidence of selection (Table S1 and Fig. 1). To the best of our knowledge, this is the first study to compile and jointly analyze all published *P. simium* samples. In contrast to previous studies that employed stringent filtering criteria for low-coverage data (*21*), we incorporated these samples using more permissive thresholds and applied a tailored suite of software and analytical approaches optimized for such data. This strategy allowed us to recover genomic variation that may have been previously overlooked. We found that the simian *P. vivax* samples from Colombia were more closely related to the Asian human *P. vivax*. The simian *P. vivax* sample in Brazil, which displayed a hybrid genetic ancestry between *P. simium* and *P. vivax*, indicated ongoing genetic exchange between these species. These results suggest the existence of an ongoing reverse zoonosis (*i.e.*, *P. vivax* transfer from humans to Neotropical NHPs), highlighting the risks of novel zoonotic malaria outbreaks and potentially affecting the health of NHPs. Furthermore, this is the first study, to our knowledge, to investigate and estimate the timing of the host jump leading to *P. simium*. We then demonstrated that *P. simium* likely originated from a transfer of human *P. vivax* to Brazilian NHPs ∼100-200 years ago, coinciding with the migration of human *P. vivax* from Mexico to this region. Lastly, we detected signals of positive selection in genes involved in interactions with primate hosts and mosquito vectors. Our results underline the importance of considering NHP as potential reservoirs of human malaria. This is crucial to anticipate potential emerging outbreaks and more globally to address human malaria public health issues but also the conservation of NHPs, whose populations might be threatened by the disease transmission.

**Figure 1.**
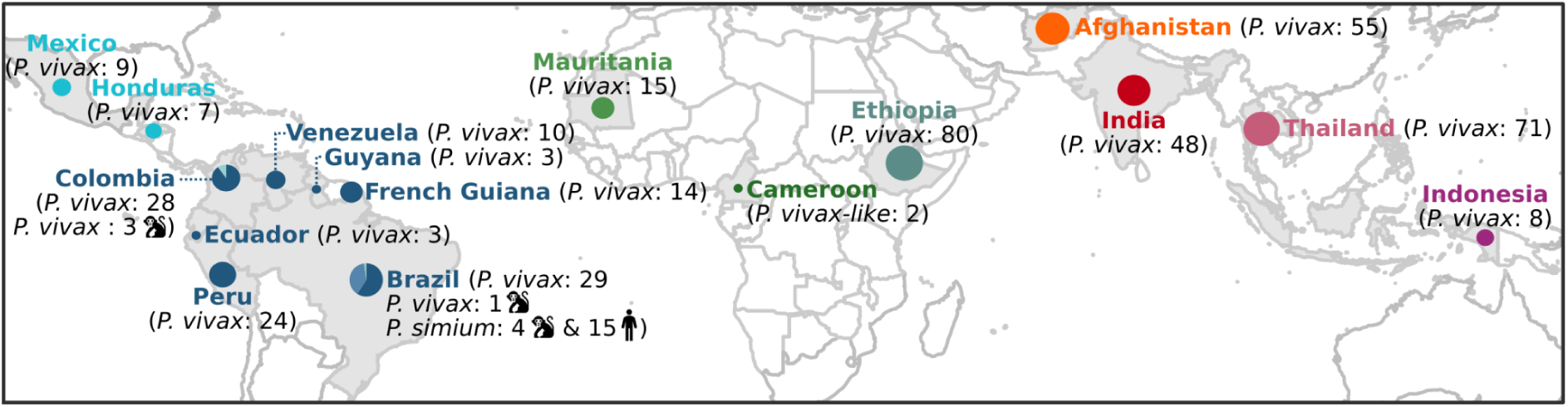
Geographic origins of the 408 *P. vivax* isolates, 19 *P. simium* isolates, and two African great apes *P. vivax-like* isolates. The number of samples for each species is indicated between brackets and by the circle size. The monkey pictogram indicates *P. vivax* and *P. simium* isolates from non-human primates in Latin America. For *P. simium* samples, the human pictogram indicates human samples.

## Results

### Evidence of *P. vivax* infections in South American non-human primates

Among 719 samples collected from NHPs across five Latin American countries (14 Colombian, 23 Brazilian, 30 Argentinian, 280 French Guianese, and 372 Costa Rican), 23 (3 from Colombia, 14 from French Guiana and 6 from Brazil) were positive for *Plasmodium* using the cytochrome-b based PCR assay (*31*) (Table S2). These 23 samples derived from various biological materials (stool, liver, mixed liver and kidney and blood) collected from five NHP species (*Alouatta macconnelli*, *Saguinus midas*, *Ateles hybridus*, *Alouatta clamitans*, and *Cebus versicolor*) (see Table S1). The sequence analysis revealed that they were all infected with *P. vivax* or *P. simium* and were all sent for whole genome sequencing (WGS), with all Colombian samples as already PCR positive according to Rondón *et al*. (*17*). Four samples, three from feces and one from blood, provided sufficient mitochondrial DNA (mtDNA) sequencing data to distinguish between *P. simium* and *P. vivax* (see Material and Methods), and were integrated with WGS data on 405 *P. vivax* and 29 *P. simium* genomes from previous studies to create a comprehensive dataset (Table S1, Fig. S1). Among the 33 *P. simium* or undetermined (*P. vivax* or *P. simium)* samples, 3 from Colombia (newly sequenced Col_164, Col_68 and Col_86) and one from Brazil (P160 from de Oliveira *et al*. (*21*)) displayed mtDNA SNPs specific to *P. vivax* (*5*, *19*, *32*), thus indicating that *P. vivax* circulates among South American NHPs.

### Genetic relationships among the *P. vivax* from Neotropical non-human primates, *P. simium,* and worldwide human *P. vivax* populations

After filtering out multi-strain infections (i.e., multiple *P. vivax* or *P. simium* genotypes) and excessive relatedness among isolates (see Materials and Methods), the final dataset included: (i) 404 human *P. vivax* samples from 15 countries; (ii) four *P. vivax* from NHPs (two from Colombian *A. hybridus*, one from Colombian *A. seniculus* and one from Brazilian *Callicebus nigrifrons*), and (iii) 19 *P. simium* isolates (n=15 from *Homo sapiens* and n=4 from *Alouatta guariba*, all from Brazil). Two additional *P. vivax*-like genomes from Cameroon chimpanzees (*Pan troglodytes ellioti*) served as outgroups (Fig. 1). From this dataset, we created two datasets with different filtering strategies to accommodate the requirements of different population genetic analyses (Fig. S1). The first dataset (*dataset_GL*) was composed of 1,286,079 SNPs and all 429 genomes, with lenient filters and accounting for genotype uncertainty through genotype likelihood analysis using the ANGSD software ecosystem (*33*). This was necessary to handle average sequencing depth heterogeneity (mean: 88.86X, median: 63.6X, with a standard-deviation of 117.27 [min: 2.5X and max: 1,142.5X]), especially for the *P. vivax* isolates from Colombian NHPs, which had low genome coverage (≥1X for 4.2% for the genome of Col_164, 11.5% for Col_68, and 8.9% for Col_86). The second dataset (*dataset_HF)* relied on more classic SNP calling using a stringent filtering procedure (see Materials and Methods). It consisted of 819,652 SNPs for 426 samples, excluding the three Colombian *P. vivax* isolates from NHPs due to their very low sequencing depth (less than 15% of genome ≥ 1X) and the large amount of missing data (>90%) (Fig. S1, Table S1 and Material and Methods). In this dataset, *P. simium* samples exhibited an average coverage of 408,268 SNPs (median: 376,932 SNPs; standard deviation: 236,902.58; range: 67,372–809,449 SNPs). In comparison, *P. vivax* samples from humans had an average SNP coverage of 703,196 (median: 769,701 SNPs; standard deviation: 152,728.37; range: 636–819,247 SNPs). The American *P. vivax* sample isolated from a monkey (sample P160) was covered by 38,682 SNPs.

Using these datasets, we first explored the genetic relationships of the simian *P. vivax* and *P. simium* isolates within the global genetic diversity of human *P. vivax* populations using multiple approaches. First, we performed a principal component analysis (PCA) and a model-based individual ancestry analysis using *dataset_GL* and *PCAngsd* (*34*) to assess the population structure. We also visualized genetic relationships among isolates using a maximum likelihood (ML) phylogenetic tree based on *dataset_HF* with IQ-TREE (*35*). The results were congruent (Fig. 2) and revealed four distinct genetic clusters of human *P. vivax*, consistent with previous studies (*36–38*): (1) East and Southeast Asia, (2) Africa, (3) Central Asia and South Asia, and (4) Latin America.

**Figure 2.**
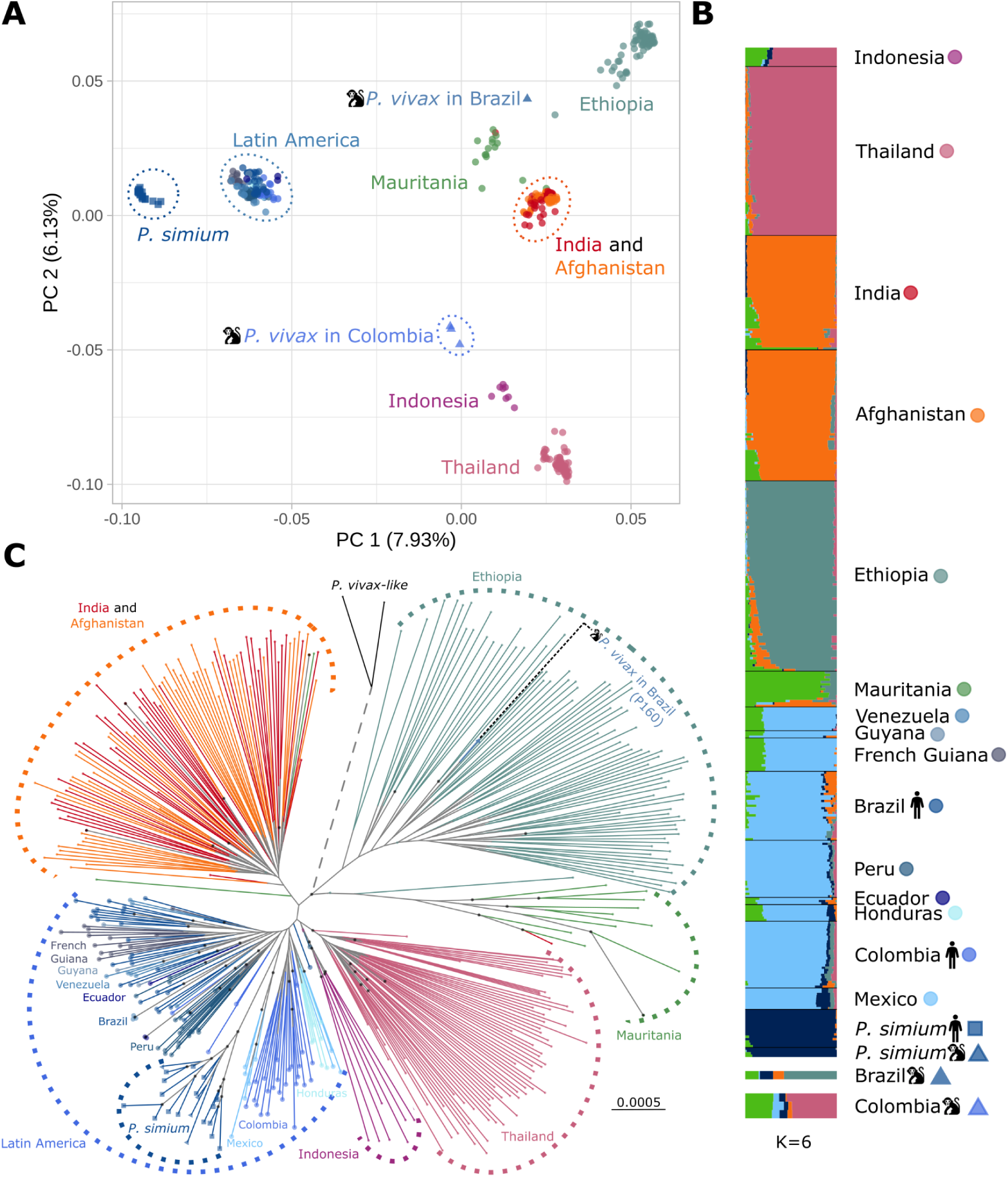
*P. vivax* and *P. simium* genetic structure. **(A)** Plot of the principal component (PC) analysis of 408 *P. vivax* and 19 *P. simium* strains showing the first and second PCs based on the genotype likelihood of 247,890 unlinked SNPs. **(B)** Individual genetic ancestry assuming K=6 genetic clusters estimated using *PCAngsd* (see Fig. S2 for the other K values). The monkey pictogram indicates the *P. vivax* samples from South American non-human primates (NHPs). **(C)** Maximum likelihood phylogenetic tree of the 404 *P. vivax*, 19 *P. simium* and 2 *P. vivax-like* individuals. *P. simium* and *P. vivax* from NHPs are indicated by triangles. The *P. vivax* isolate from a Brazilian NHP is among the Ethiopian *P. vivax* isolates. The three *P. vivax* samples from Colombian NHPs could not be included in this analysis because of the high number of missing data. The tree includes two *P. vivax-like* strains used as outgroups. Note that the length of the outgroup branch (dotted lines) has been truncated. Black dots at nodes indicate highly supported nodes (both SH-aLRT ≥ 80% and UFboot ≥ 95%, following the thresholds by Guindon *et al.* (*39*) and Minh *et al.* (*40*)).

The four *P. vivax* isolates from NHPs did not cluster with *P. simium* isolates in the PCA (Fig. 2A). The three Colombian simian *P. vivax* isolates were closely related to South-East Asian *P. vivax* isolates, while the Brazilian isolate (P160) grouped together with African *P. vivax*, particularly Ethiopian samples (Fig. 2A). The Colombian samples displayed admixed genetic ancestry (Fig. 2B), with contributions from South-East Asian and West African *P. vivax* populations. Conversely, the Brazilian sample shared genetic ancestry with *P. simium* and predominantly with Ethiopian *P. vivax* (Fig. 2B). This isolate clustered within the Ethiopian diversity in the ML tree, although the node support was not high (SH-aLRT = 95% and UFboot = 91%, using thresholds established by Guindon *et al.* (*39*) and Minh *et al.* (*40*)) (Fig. 2C).

In the PCA plot (Fig. 2A), *P. simium* strains, isolated from NHPs and humans, formed a distinct cluster, separated from human *P. vivax* along the first principal component (PC 1). *Plasmodium simium* isolates were genetically closer to American *P. vivax* populations than to other populations from the rest of the world. The genetic ancestry plots further supported this result. Indeed, *P. simium* clustered with the American *P. vivax* when testing a number of clusters (K) ranging from 2 to 4 (Fig. S2). Only with K=5, *P. simium* started to differentiate (Fig. S2), although it still shared genetic ancestry with the Mexican *P. vivax* (dark blue in Fig. 2B). The ML tree supported this result: *P. simium* branched within the genetic diversity of Mexican *P. vivax*, despite limited node support (SH-aLRT = 100% and UFboot = 80%, using thresholds established by Guindon *et al.* (*39*) and Minh *et al.* (*40*)).

Although *P. simium* formed a well-defined group distinct from the human *P. vivax* (Fig. 2), the strains originated from various regions in Brazil and from different host species (humans and *A. guariba*) (Fig. 3A). Therefore, we investigated whether there was a genetic sub-structuring within *P. simium*, independently of *P. vivax,* according to the host species and geography. The genetic ancestry analysis with *PCAngsd* (*34*) identified two genetic clusters within *P. simium* Fig. 3B, and Fig. S3). This separation was also evident in the first principal component of the PCA (Fig. 3C). The Rio de Janeiro (RJ) cluster (n=13) comprised *P. simium* samples from humans in the states of Rio de Janeiro and Espírito Santo, as well as in the municipality of Peruíbe, in the State of São Paulo. Additionally, *P. simium* samples from *A. guariba* were included from Rio de Janeiro and São Paulo (Fig. 3B). Within this cluster, the three samples from Rio de Janeiro, isolated from NHPs and humans, were distinct from the other samples in the RJ cluster along the second principal component of the PCA (Fig. 3C). In contrast, the São Paulo (SP) cluster (n=6), located further south, included only human samples from the municipalities of Maresias, Riacho Grande, Juquitiba, Registro, and Iporanga, all within the state of São Paulo. Additionally, a sample from the RJ cluster (originating from Peruíbe) appears to exhibit admixture with the SP cluster, likely due to its geographical proximity (Fig. 3B).

**Figure 3.**
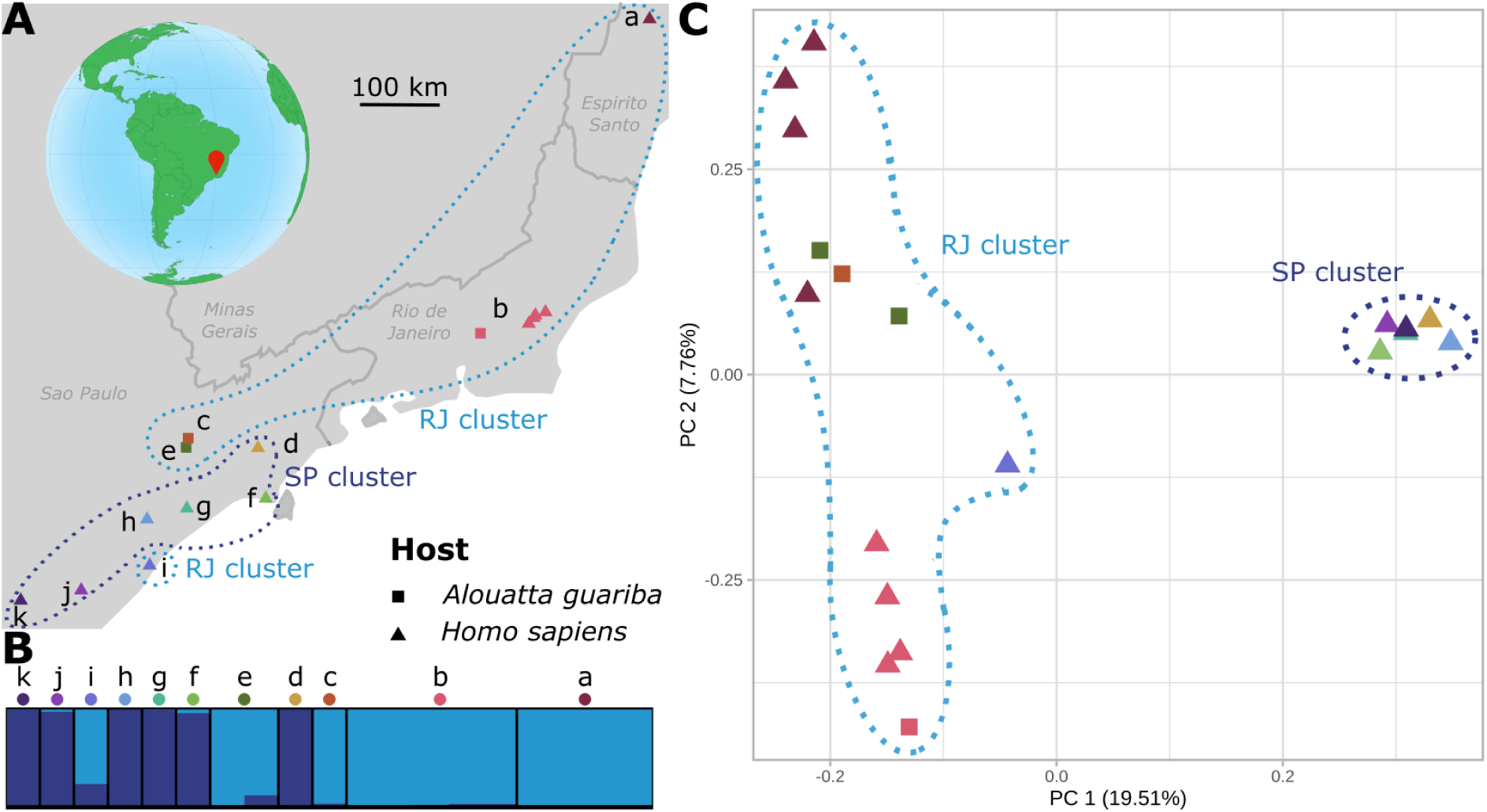
Geographic origins and genetic structure of the Brazilian *P. simium* isolates. **(A)** Geographic origins of the 19 *P. simium* isolates. The squares and triangles represent the host species (non-human primates and humans, respectively), and the colors correspond to the region of origin. The SP cluster is outlined in dark blue, the RJ cluster in light blue.The borders and names of the Brazilian states are indicated in dark gray. On the globe, the red pointer indicates the region of Brazil from which the samples originate. Each site is designated by a letter: (a) Espírito Santo (n=4), (b) Rio de Janeiro (n=5), (c) São Paulo, Mairipor (n=1), (d) Paraibuna (n=1), (e) São Paulo, Cantareira State Park (n=2), (f) Maresias (n=1), (g) Riacho Grande (n=1), (h) Juquitiba (n=1), (i) Peruíbe (n=1), (j) Registro (n=1), (k) Iporanga (n=1). **(B)** Individual genetic ancestry assuming K=2 genetic clusters estimated using *PCAngsd* (see Fig. S3 for the other K values). Each site is designated by a letter: (a) Espírito Santo (n=4), (b) Rio de Janeiro (n=5), (c) São Paulo, Mairipor (n=1), (d) Paraibuna (n=1), (e) São Paulo, Cantareira State Park (n=2), (f) Maresias (n=1), (g) Riacho Grande (n=1), (h) Juquitiba (n=1), (i) Peruíbe (n=1), (j) Registro (n=1), (k) Iporanga (n=1). **(C)** Plot of the principal component (PC) analysis of 19 *P. simium* strains showing the first and second PCs based on the genotype likelihood of 44,911 unlinked SNPs. The shape (square and triangle) represents the host species (non-human primates and humans), and the color the region of origin. The SP cluster is outlined in dark blue, the RJ cluster in light blue.

### Evidence of admixture between *P. simium* and *P. vivax*

We next investigated the role of admixture in shaping the genetic makeup of different populations and samples. We estimated and visualized the pairwise identity-by-descent (IBD) as a network depicting recent common ancestry among samples (*41*, *42*) (see the Materials and Methods). We also constructed population graphs using *AdmixtureBayes* and *TreeMix* (*43*, *44*), to draw information on population branching order and potential admixture events (*i.e.* reticulation). We complemented these analyses with the *f_4_-statistics* which formally test for admixture among populations (*45*, *46*). All these analyses were based on the *dataset_HF* (Fig. S1).

As observed in the population structure analyses (Fig. 2), the pairwise IBD network showed the same four main genetic clusters identified in *P. vivax*: Southeast Asia, Central and South Asia, East Africa, and Latin America. In the IBD network, all *P. simium* isolates formed a single coherent and genetically distinct cluster, closely related to Latin American *P. vivax* (Fig. 4A). Conversely, the Brazilian simian *P. vivax* (P160) isolate was genetically distinct, showing strong connections with Ethiopian *P.vivax* and *P. simium* isolates, as observed in previous analyses (Fig. 2).

**Figure 4.**
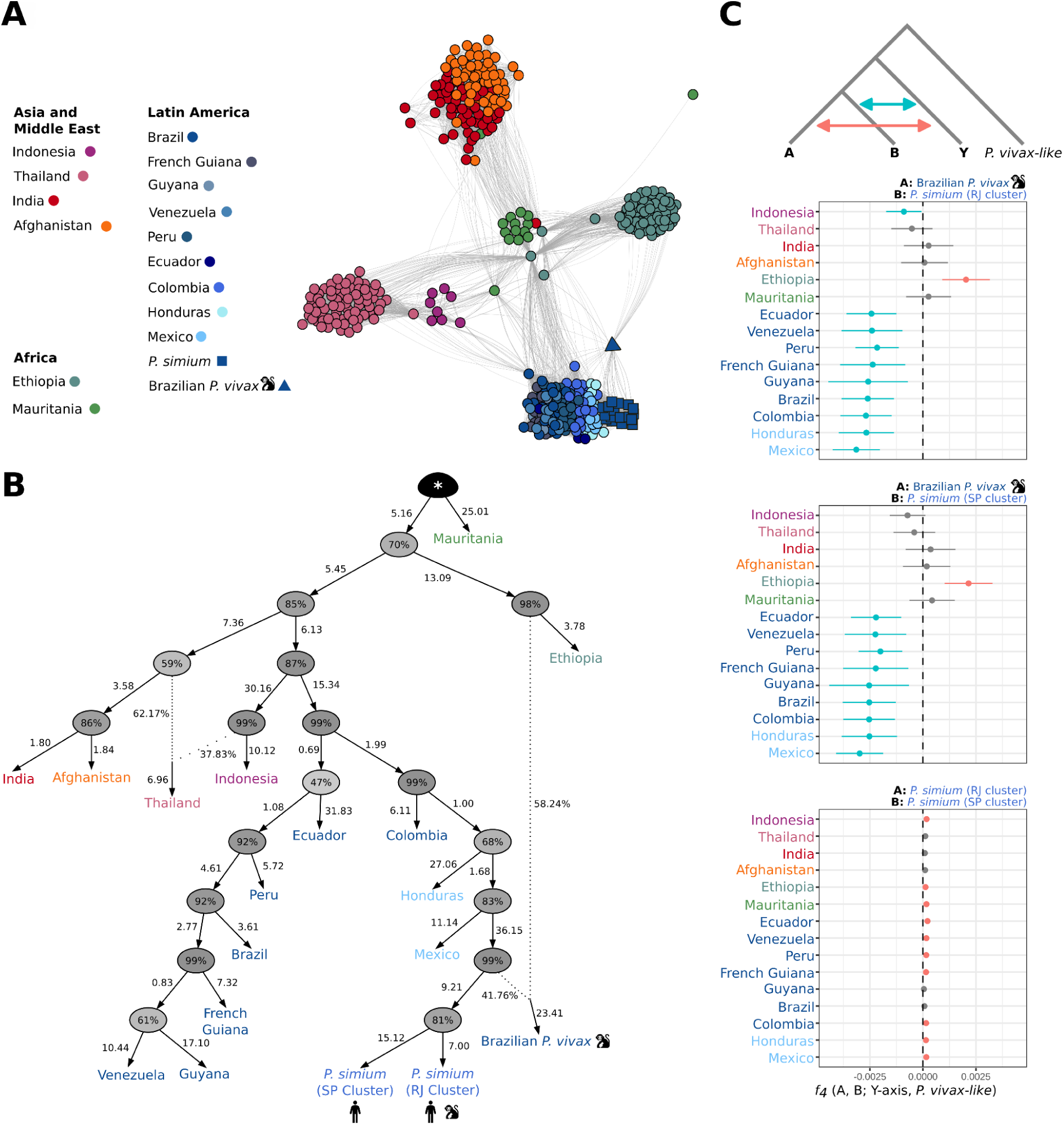
Population genetic relationships and admixture between *P. vivax* and *P. simium.* **(A)** Network visualization of the identity-by-descent (IBD) relationships among *P. vivax* and *P. simium* sample pairs. Edges represent the IBD values between each sample pair. Only edges with IBD values > 5% are displayed. **(B)** Admixture population graph topology of *P. vivax* and *P. simium* populations. The highest posterior probability was estimated with *AdmixtureBayes*(*43*), and rooted with the two *P. vivax-like* genomes (*). The branch length (measured in drift units) indicates the genetic divergence between populations, multiplied by 100. Percentages at nodes indicate the posterior probability that the true graph has a node with the same descendants. For each admixture event (indicated by the dotted lines), the percentages show the admixture proportions from each donor. **(C)** Admixture tests based on the *f_4_-statistics* for *P. simium* clusters and the Brazilian *P. vivax* sample from a non-human primate with *P. vivax* populations. This statistic assesses the excess of shared alleles between populations. In the form *f_4_*(A, B; Y-axis, P. vivax-like), values not significantly different from 0 (gray) indicate no gene flow between populations A or B and the Y-axis population. Positive values (red) suggest gene flow between population A and the Y-axis population, and negative values (blue) indicate gene flow between population B and the Y-axis population. The monkey pictogram indicates the *P. vivax* isolate from Brazilian non-human primate.

The *AdmixtureBayes* analysis identified a population graph with two admixture events as the best-fitting solution for our data (Fig. 4B and Fig. S4). *P. simium* isolates formed a well-supported monophyletic group (posterior support >80%) that branched off from Mexican *P. vivax* (Fig. 4B). The first reticulation in the *AdmixtureBayes* population graph identified Thailand *P. vivax* as an admixed population between Indonesian populations and the common ancestor of the Indian and Afghan populations. This reticulation was not visible in the pairwise IBD network, which only showed recent ancestry, and in *TreeMix*, which identified only one optimal migration edge (Fig. S5). The only reticulation detected by *TreeMix*, also present in the *AdmixtureBayes* population graph, suggested that the Brazilian simian *P. vivax* isolate resulted from an admixture event between East African *P. vivax* and *P. simium*, with ∼40% of its ancestry from *P. simium* (Fig. 4B and Fig. S5). These results were supported by the *f_4_*-statistic tests (Fig. 4C), confirming that this simian *P. vivax* sample was genetically admixed and closer to Ethiopian *P. vivax* than to the *P. simium* cluster.

### *Plasmodium simium* demographic history

We further investigated *P. simium* demographic history, potential founder effect, and isolation dynamics. We first computed the genome-wide distribution of the nucleotide diversity (π) in windows along the core genome of populations with at least 10 samples. *P. simium* exhibited significantly lower π values than *P. vivax* (*p-value* <0.001, Wilcoxon signed-rank test with Bonferroni correction, Fig. S6). Then, we explored the changes in effective population size (*N_e_*) over time, estimated using coalescence rates with the *Relate* software (*47*). In agreement with the π values (S6 Fig.), both *P. vivax* and *P. simium* populations worldwide exhibited a steady decline in *N_e_* starting ∼100,000 generations ago or ∼20,000 years ago (assuming a generation time of 5.5 generations/year and a mutation rate of 6.43 x 10^-9^ mutations/site/generation (*36*)), followed by a recent moderate expansion (Fig. 5A). Notably, the two *P. simium* clusters (RJ and SP) displayed lower *N_e_* values than the *P. vivax* populations, although we did not detect any strong signal of bottleneck associated with a potential founder effect.

**Figure 5.**
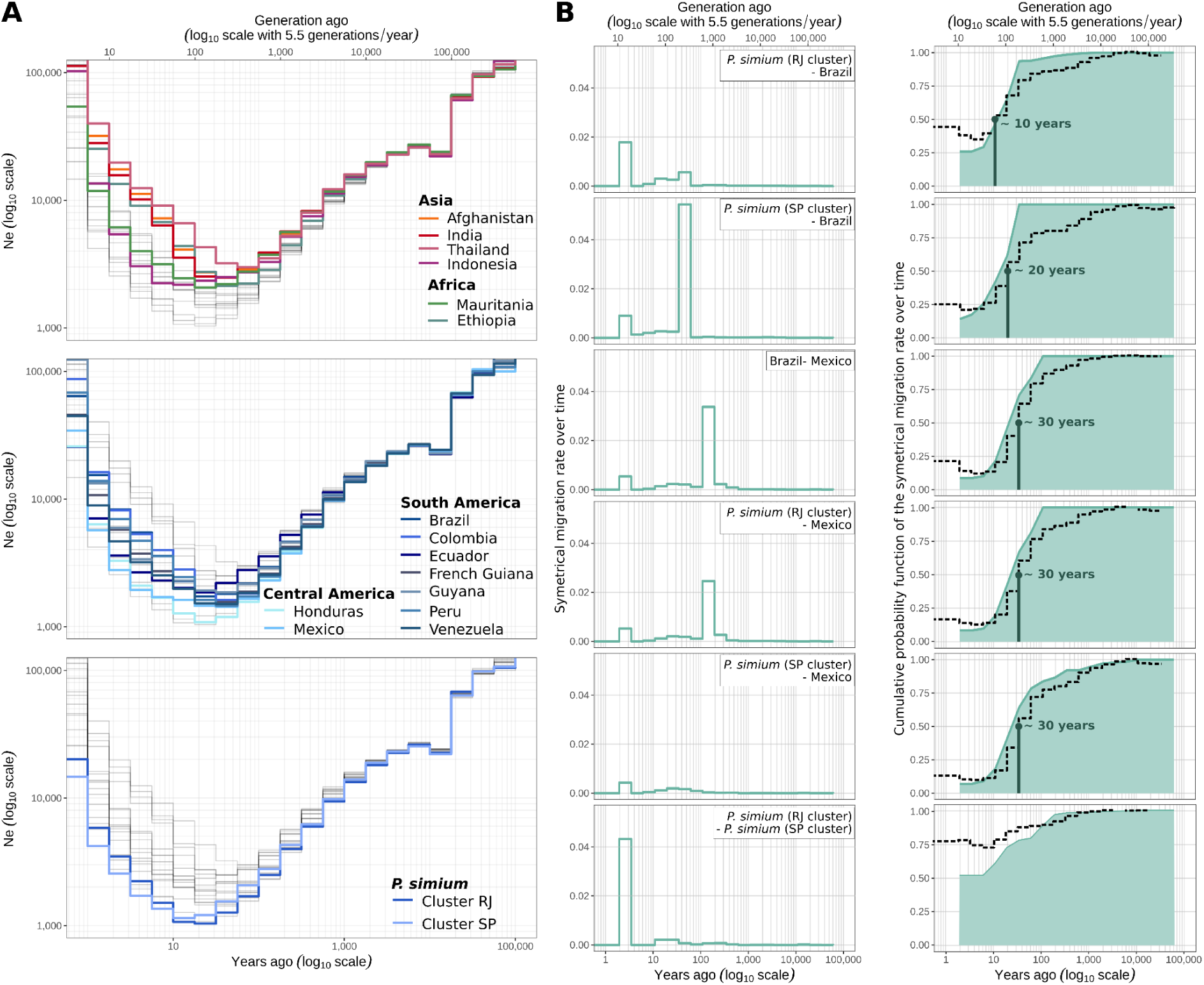
Coalescent-based inference of the demographic history and migration rates of *P. vivax* and *P. simium* populations. **(A)** The variation in effective population (*Ne*) size was estimated using *Relate* (axes are log_10_ transformed). **(B)** Timing and dynamics of the separation between the indicated groups estimated using *Relate* coalescence rates and *MSMC-IM*. On the left, the time-dependent-symmetric migration rate (x-axes are log_10_ transformed). On the right, the cumulative distribution probabilities of the migration rates in function of time. This is an estimate of the proportion of ancestry already merged at time *t*, and represents the proportions of gene flow over time (x axes are log_10_ transformed). Values close to 0 indicate complete isolation between groups, while 1 shows a complete mixture as one population. Dashed lines indicate the relative cross-coalescence rate. The divergence time (when the cross-coalescent rate drops below 50%) is specified in each panel.

To explore the isolation and migration history of *P. simium* from *P. vivax* in Latin America, we used Multiple Sequentially Markovian Coalescent-Isolation Migration program (MSMC-IM) (*48*) to estimate the isolation dynamics and the migration rate changes over time. Our results, based on the cross-coalescent rates (CCR) between *P. simium* and American *P. vivax* populations, suggested a relatively recent isolation of *P. simium* from *P. vivax*. Based on the point at which the CCR dropped below 50% as suggested by Wang *et al.* (*48*), we could estimate that in all pairwise comparisons between *P. simium* and American *P. vivax*, split times occurred between 10 and 30 years ago (Fig. 5). The only exception was between the *P. simium* clusters, where CCR values remained ∼75%, indicating that genetic exchange was still ongoing (Fig. 5B, bottom).

Migration rate estimations (Fig. 5B) suggested that the oldest migration events occurred 100-200 years ago between the Mexican and Brazilian *P. vivax* populations, and between Mexican *P. vivax* and *P. simium* from the RJ cluster. The estimated divergence times were ∼30 years ago for both the Mexican-Brazilian *P. vivax* and Mexican *P. vivax*-*P. simium* RJ cluster. Similarly, the divergence time between the *P. simium* SP cluster and Mexican *P. vivax* was ∼30 years ago, with minimal genetic exchange over the entire period (Fig. 5B). Conversely, the *P. simium* SP cluster and Brazilian *P. vivax* populations have exchanged genetic material more recently (30-60 years ago), with divergence estimated at ∼20 years ago. On the other hand, the RJ cluster showed limited recent exchanges with Brazilian *P. vivax*, and diverged ∼10 years ago. Importantly, although the cumulative migration probability (*Mt*), which is analogous to the CCR (*48*, *49*), was close to zero for some comparisons between *P. vivax* and *P. simium*, none of them reached zero. This suggests that genetic exchanges may still be possible, even at a low rate (Fig. 5B).

### Genetic evidences of *P. simium* adaptation to new hosts

The host jump from humans to NHPs likely exerted strong selective pressure on *P. simium*, potentially leaving detectable signals of positive selection in its genome. Therefore, we first used haplotype-based tests within (*iHS* (*50*)) and between (*XP-EHH,* and *Rsb* (*51*, *52*)) populations to identify signals of positive selection specific to a target population. These tests use haplotype lengths variations (*i.e.,* linkage disequilibrium, LD) to detect recent or ongoing selective events (*50–52*). Given the limited sample size (n=19), we treated the two *P. simium* clusters as a single group and compared it to the Brazilian *P. vivax* population as the reference population for *XP-EHH* and *Rsb* statistics. The datasets derived from *dataset_HF,* following the determination of ancestral and derived alleles with *P. vivax-like* genomes, included 10,304 SNPs for the 19 *P. simium* samples for the *iHS* test, and 378,441 SNPs for 48 samples (n=19 *P. simium* and n=29 Brazilian *P.vivax*) for the *XP-EHH* and *Rsb* analyses (see Material and Methods for more details). The within-population *iHS* test identified a single significant SNP within the coding sequence (CDS) of the *PVP01_1139100* gene, an orthologue of a *Plasmodium* protein of unknown function (Fig. S7). The *XP-EHH* and *Rsb* analyses identified two significant SNPs in the CDS of the *PTEX150* gene (Fig. 6A, Fig. S7 and Table S3), which encodes a protein involved in protein transport across the parasitophorous vacuolar membrane during the *Plasmodium* erythrocytic cycle (*53*, *54*).

**Figure 6.**
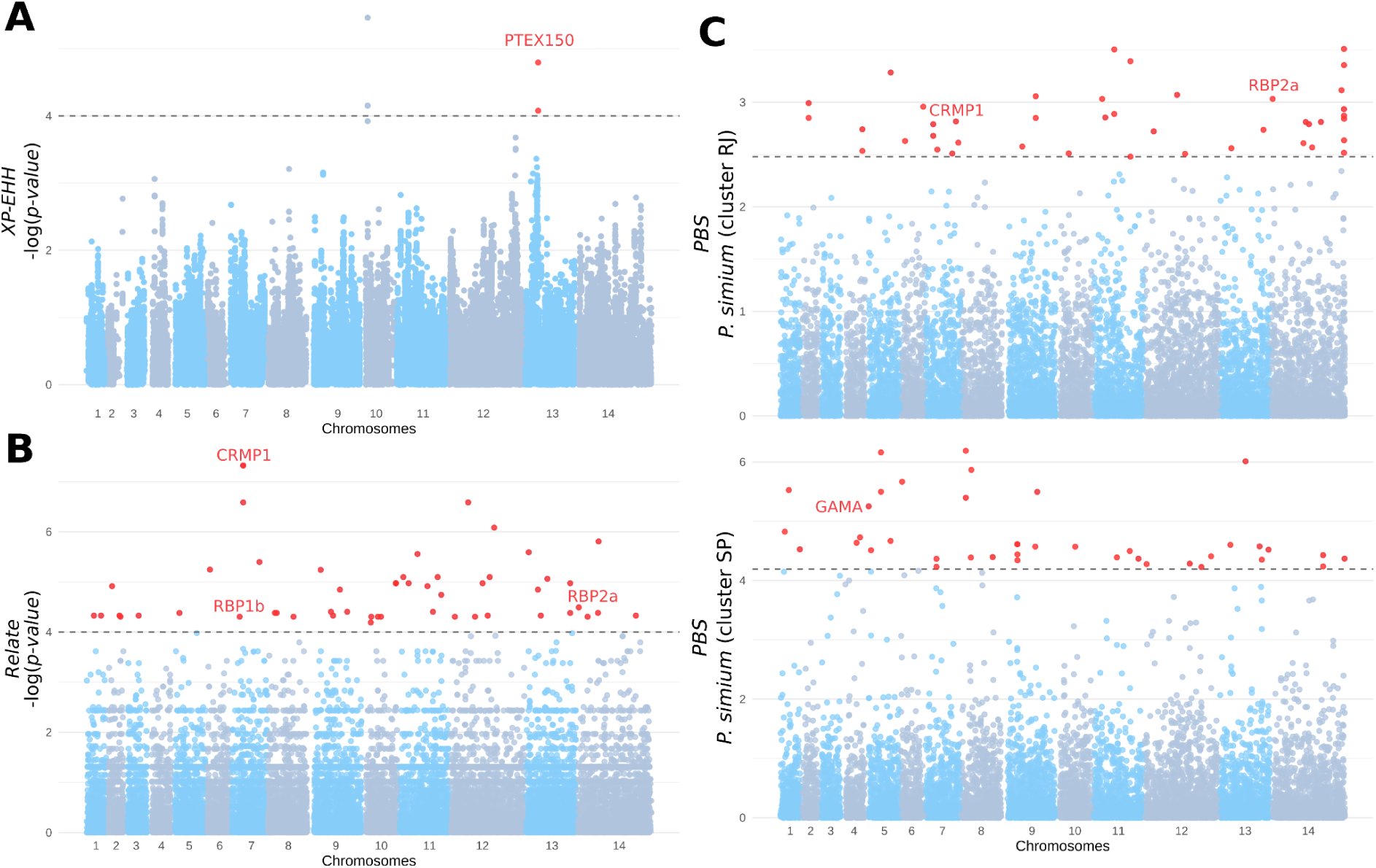
Evidence of selective sweeps in *P. simium.* **(A)** Manhattan plot showing the haplotype-based *XP-EHH* statistics between pairs of populations. The dotted line represents the significance threshold value at -log(*p-value*) = 4. Red points are the SNPs identified as affected by a selective sweep in *P. simium* (negative *XP-EHH* values). **(B)** Manhattan plot showing the *Relate* results. The dotted line represents the threshold significance value -log(*p-value*) = 4. The red points are the SNPs identified as potentially affected by positive selection in *P. simium*, on the basis of their higher-than-expected coalescence rate compared with the rest of the genome. **(C)** Manhattan plots showing the *PBS* values, an *F_ST_*-type statistic that captures a difference in allele frequencies compared with two reference populations, for each *P. simium* cluster, with *P. vivax* Brazilian and Colombian samples as outgroup populations. For the *PBS* scores, all values in red represent the top 0.1% of the PBS values, suggesting higher allelic frequency differentiation compared with the reference populations, and thus indicating significant evidence of positive selection that affects these SNPs. CRMP1, cysteine repeat modular protein 1; RBP1b, reticulocyte binding protein 1b; RBP2a, reticulocyte binding protein 2a; GAMA, GPI-anchored micronemal antigen.

The haplotype-based tests may miss other genomic regions because recombination can break down LD patterns. Therefore, we used *Relate* (*47*) to detect positive selection by identifying increased coalescence rates in selected regions compared with the rest of the genome. By analyzing 26,565 SNPs in the 19 *P. simium* samples, we identified 49 SNPs with significant signals of selection in the CDS of 14 genes (Table S3). Notably, three of these genes are involved in the interactions of the parasite with primate hosts (*PvRBP2a* and *PvRBP1b* (*55–57*)) and mosquitoes (*PvCMRP1* (*58–60*)).

We also investigated cluster-specific evidence of adaptation in *P. simium* by calculating the population branch statistic (*PBS*), an *F_ST_*-like statistic, by comparing each *P. simium* cluster with two American *P. vivax* populations (Colombia and Brazil). Outlier values (top 0.1% of the 41,338 1-kb sliding windows for the RJ cluster and of the 39,799 1-kb sliding windows for the SP cluster) indicated significant differentiation of the *P. simium* clusters compared with American *P. vivax*. We identified 43 outlier windows in the RJ cluster and 41 in the SP cluster, corresponding to positive selection in the CDS of 32 genes (RJ cluster) and 31 genes (SP cluster) (Table S3). Four genes showed signs of positive selection in both clusters (*PVP01_0525300*, *PVP01_0931400*, *SMC1* and *PVP01_1469500*). *SMC1* (Structural Maintenance of Chromosomes protein 1) is involved in the structural maintenance of chromosomes in eukaryotes (*61*, *62*). Conversely, the functions of the other three genes are unknown. In the RJ cluster, two genes with evidence of selection have already been detected with *Relate*: *PvRBP2a* and *PvCMRP1.* These play a direct role in the interactions with the primate hosts and with mosquitoes, respectively. The SP cluster showed selection signals in *PvGAMA*, a gene involved in binding to human erythrocytes (*63*).

In conclusion, we identified 73 genes under positive selection in *P. simium*, including several implicated in interactions with both mosquito vectors and primate hosts.

## Discussion

Historically, natural *P. vivax* or *P. simium* infections in NHPs (*A. guariba* and *A. caraya*) in Brazil were detected with antigenic tests (*64*) and/or PCR assay (*16*), although the two species were often indistinguishable from one another. *Plasmodium simium* was long thought to circulate exclusively in Brazil (*5*, *64*, *65*). Recently, this assumption was challenged by studies that reported natural infections in Colombian and Costa Rican NHPs, attributed to *P. vivax* or *P. simium* (*17*, *18*). This questioned the natural range of *P. simium* in Neotropical NHPs and its geographic distribution. Here, after screening 719 NHPs samples from five different Latin American countries, *P. simium* seemed restricted to Brazil. For the first time to our knowledge, we identified natural infections by *P. vivax* in two American NHP species (*A. hybridus* and *A. seniculus*). The analysis of the published *P. simium* genomes allowed the identification of another *P. vivax* isolate in a NHP from Brazil (sample “P160”). These findings suggest ongoing host switches from humans to American NHPs, highlighting the reverse zoonotic potential of these parasites as already suggested (*21*).

The three Colombian *P. vivax* isolated from NHPs were genetically closer to South-East Asian *P. vivax* than to the American strains (Fig. 2). This pattern appears to be specific to simian *P. vivax* and is not observed in American isolates sampled from humans (*36–38*, *66*). This genetic relationship may reflect historical human migrations from Asia or Oceania to Latin America (*67*), but it could be also linked to international animal trade between these regions (*68*), including the introduction of Asian NHPs transported to Brazil for experimental purposes (*69*). These findings suggest a potential re-invasion of Asian human *P. vivax* into American NHPs. More simian *P. vivax* sampling in Colombia with greater sequencing depth is needed to clarify this evolutionary history.

In this study, the *P. vivax* sample P160 from a Brazilian NHP was reanalyzed using methods optimized for low-coverage data and less stringent filtering than in previous work (*21*), enabling us to recover genomic variation that had been previously overlooked. This analysis revealed traces of admixture between *P. simium* and Ethiopian *P. vivax* (Fig. 2B and Fig. 3), consistent with prior observations (*21*). The recent admixture detected in this sample demonstrates ongoing recombination between these species. The genetic proximity or even lack of distinct polymorphisms on some markers between *P. simium* and *P. vivax* had already raised questions about their classification as separate species (*21*, *70*). This is not surprising when considering that the host jump occurred only ∼80-100 years ago, and that we detected ongoing gene flow between these species (Fig. 5B). Interestingly, the P160 sample showed a mixture with Ethiopian rather than American *P. vivax*, which is unexpected given its collection area. The lack of literature explaining this Ethiopian ancestry underscores the need of more sampling to better understand the history of this sample.

The discovery of natural *P. vivax* infections in Colombian NHPs suggests that NHPs could act as anthropo-zoonotic reservoirs, thus complicating malaria elimination efforts across the continent, but also affect the survival of NHPs infected. *Plasmodium vivax* remarkable adaptability in changing hosts contrasts with *Plasmodium* species of the subgenus *Laverania* (including *P. falciparum*, the deadliest human malaria agent) where host transfer is rarely observed (*6*). Ongoing genetic exchanges between *P. vivax* and *P. simium* indicate porous species boundaries and suggest that recombination still occurs. These results are of major importance for malaria epidemiology and underscore the role of NHPs as potential malaria reservoirs. Therefore, it is crucial to consider NHP hosts in strategies to control and potentially eliminate malaria in Latin America. These strategies must also consider the health and conservation of already threatened NHPs (*71*), which have previously been killed by humans to prevent yellow fever transmission in Brazil (*72*). Furthermore, the killing of NHPs may have negative impacts on public human health in both the short and long term (*72*).

The evolutionary processes underlying *P. simium* host shift and adaptation have not been fully explored so far (but see de Oliveira *et al.* (*21*)). Our PCA, ancestry plots, ML phylogenetic tree (Fig. 2), IBD networks, and population graphs (Fig. 4A-B) consistently showed a genetic structure among *P. vivax* populations that is in line with previous findings (*36–38*). *P. simium* formed a unique cluster closely related to American *P. vivax*, particularly Mexican strains, as supported by its branching within the Mexican diversity in the ML tree and population graph (Fig. 2C, Fig. 4B, Fig. S5), as previously reported (*20*, *21*). Our results, supported by the ancestry plot (Fig. 2B), the population graph, and the *f_4_*-statistics (Fig. 4), showed no genetic evidence of any Asian contribution to *P. simium* genetic ancestry, unlike some early studies based on mitochondrial DNA (*24*, *25*). This supports an American origin for *P. simium*, closely linked to Mexican *P. vivax*.

A closer look at the genetic structure of *P. simium* isolates revealed two distinct genetic clusters (Fig. 3): RJ and SP. The SP cluster contains only human *P. simium* samples, whereas the RJ cluster included *A. guariba* and human samples. This is the first time this structure has been observed in *P. simium*, possibly thanks to the integration of different datasets in this study. However, the structure appears more complex. Indeed, the PCA results indicate that samples from Rio de Janeiro city are genetically distinct within the RJ cluster, while the Peruíbe sample is geographically nearer to the SP cluster, but its genetic ancestry is mainly from the RJ cluster. More genomic data across regions and host species are needed to clarify these patterns. For example, since the states of São Paulo and Rio de Janeiro are highly connected by public transports, the movement of infected humans can not be excluded to explain the observed pattern.

The close genetic similarity between human *P. vivax* and NHP *P. simium* suggests recent host transfer(s), as previously proposed (*14*, *20*, *21*), although no precise split times was estimated. Our coalescent-based analyses estimate that *P. simium* diverged from American *P. vivax* in the last 30 years (Fig. 5B). Our analyses also suggest that significant migration occurred ∼100-200 years ago between Mexican and Brazilian *P. vivax* and between the *P. simium* RJ cluster and Mexican *P. vivax* (Fig. 5B). These results suggest a first host jump might have occured ∼100-200 years ago when Mexican *P. vivax* migrated to Brazil. This timing coincides with the migration of several thousand confederates from the Southern United States to Latin America, including Mexico and Brazil, at the end of the American Civil War (*73*, *74*). It is plausible that *P. vivax* strains from the Southern United States, potentially genetically close to Mexican strains due to their geographical proximity, contributed to the origin of the RJ cluster in *P. simium*. To confirm this hypothesis Southern United States samples should be sequenced to assess their genetic connection to Mexican strains and their potential role in the emergence of *P. simium* in Brazil.

The *P. simium* SP cluster did not show any evidence of genetic exchanges with Mexican *P. vivax*, but exhibited gene flow with Brazilian *P. vivax* that occurred ∼30–60 years ago. The *MSMC-IM* analysis indicated ongoing gene flow between the *P. simium* clusters, with CCR values well above 50% (threshold to suggest split time following Wang *et al.* (*48*)) and close to 75%, along with the high migration rate ∼2 to 4 years ago. This suggests that the two *P. simium* populations have not fully diverged and may still exchange genes. Thus, the SP cluster origin remains unresolved. Indeed, population graphs showed that both *P. simium* clusters connected to the ancestor of Mexican *P. vivax* (Fig. 5), while the *MSMC-IM* results indicated that the SP cluster may have originated between 30 and 60 years ago from human *P. vivax* from Brazil (Fig. 6). The close genetic proximity between the *P. simium* clusters may reflect recent secondary contacts, as suggested by the recent high rate of migration (∼2-4 years ago, see Fig. 5B) and the presence of an admixed individual (Fig. 3B). Nevertheless the SP cluster small sample size (n=6) limits conclusions about its origin and evolutionary dynamics. Previous studies proposed the possibility of multiple host jumps based on the presence of polymorphisms in few mitochondrial or nuclear markers (*22*, *23*). An alternative hypothesis could be a single host jump that involved several *P. vivax* strains infecting NHPs. Therefore, we cannot rule out the possibility that the SP cluster derived from the RJ cluster and that the genetic distinction observed between clusters might be attributed to host adaptation dynamics. Specifically, the RJ cluster infects both simian and human hosts, whereas the SP cluster seems to infect exclusively humans. Larger samples are needed to determine the role of host specificity in the observed genetic divergence, but also the influence of human movement between these two clusters. Scenario-testing methods, such as *DIYABC-RF* (*75*), *dadi* (*76*) and *fastsimcoal* (*77*), could also be useful to clarify *P. simium* evolutionary history.

Importantly, the inferred divergence times may be biased by the life-history traits of *P. simium* and *P. vivax*. *Relate* and *MSMC-IM* rely on the Wright-Fisher (WF) coalescent model (Kingman’s coalescent model (*78*)) that assumes a panmictic population with non-overlapping generations and neutral evolution (*79*, *80*). However, this assumption does not hold for *P. vivax* and likely for *P. simium* because they exhibit dormancy and emerge when the host is infected with new strains (*81*, *82*). Consequently, diversity is maintained, the effect of drift is attenuated, and the effective population size may be overestimated (*83*, *84*). This could explain the absence of a detectable host-transfer bottleneck in the *P. simium* population size inferences (Fig. 5). Moreover, the Kingman’s coalescent model assumes only the coalescence of two lineages per generation (*78*). However *Plasmodium* life cycle alternates between asexual reproduction in primates and sexual reproduction in mosquitoes, potentially leading to simultaneous coalescent events (*85*). When such events are frequent the Kingman’s model underestimates the effective population sizes, although it remains accurate if these events are rare (*86*). To address these biases, we excluded clonal and related individuals (see Materials and Methods). Despite these precautions, the WF model limitations and the unique biology of *P. simium* and *P. vivax* suggest that divergence times and population size estimates should be interpreted with caution because they may not fully capture the parasite complex evolutionary dynamics. Moreover, estimates of effective population size and divergence time are influenced by the choice of mutation rate, which is inherently challenging to determine. In this study, we selected a value commonly used in the literature (*36*, *37*), but we recognize that mutation rates can vary spatially and temporally in *P. vivax*. However, we would like to emphasize that this mutation rate has previously been discussed as a reasonable approximation for *P. vivax* (*37*). All these biases could therefore explain why we find divergence dates more recent than the first descriptions of *P. simium* (*13*, *15*, *20*).

Through its evolution, *P. simium* has experienced both human-to-wild animal and wild animal-to-human transmission, making it an interesting candidate for studying the genetics basis of host adaptation. The recent host shift has likely imposed significant selection pressure on *P. simium* genomes, and our results provide evidence of such selective processes on the standing genetic variation. Specifically, we found significant signals in SNPs located in two genes (*PvRBP2a* and *PvRBP1b)* that play a crucial role in the interaction with the NHP host red blood cells (Fig. 6). These proteins, apically expressed in merozoites, facilitate *P. vivax* binding to reticulocytes (*55–57*). Previous studies identified a deletion in *P. simium PvRBP2a* CDR and divergence between *P. vivax* and *P. simium* (*20*, *21*). Our findings suggest that these selection signals may reflect linked selection near the deletion (∼1,912 nucleotides away). This is the first report of positive selection in *PvRBP1b* in *P. simium*. Given the essential role of these two genes in reticulocyte invasion (*55–57*), our results suggest that *P. simium* has adapted to its new hosts to effectively invade their reticulocytes. It would therefore be relevant to study the impact of this parasite on the health and survival of the NHPs in this region, which are already threatened (*71*). Noteworthy, within the SP cluster, which comprises *P. simium* samples that infect only humans, we discovered significant divergence in the *PvGAMA* gene compared with American human *P. vivax* (Fig. 6C). *PvGAMA* binds to human erythrocytes irrespective of their Duffy antigen status(*63*), suggesting that some *P. simium* genetic groups may have specifically adapted to human hosts. Functional studies on reticulocyte invasion in simian and human hosts based on the *PvGAMA* genotypes will be crucial to determine the gene role in host adaptation.

*P. simium* adapted not only to primate hosts but also to new mosquito vectors. Unlike *P. vivax*, *P. simium* is thought to be confined to the Atlantic coast in Brazil and transmitted primarily by anopheline mosquitoes of the *Kerteszia* subgenus, mainly *An. cruzii* and *An. bellator* (*29*, *30*). We identified selection signals in the *PvCMRP1* gene for all *P. simium* samples and in the *PvPAT* gene only for the SP cluster (Fig. 6). Previous studies showed that *PvCMRP1* underwent positive selection in Latin American *P. vivax* populations (*38*, *87*, *88*), suggesting that the signal observed in *P. simium* may reflect a similar evolutionary pressure in American *P. vivax*. Although the exact functions of *PvCMRP1* and *PvPAT* are not fully known, studies in which their orthologs were knocked out showed that they are crucial for completing the mosquito life cycle and transmission to vertebrate hosts(*58–60*).

de Oliveira *et al.* (*21*) detected selection signals in the *pvs47* gene (potentially essential for avoiding the mosquito immune system), the sequence of which differs between *P. vivax* and *P. simium* by only two amino acid substitutions (*89*). Although both SNPs are present in our dataset, our analyses did not find them to be under selection. The small *P. simium* sample size (n=19) may have limited the power of our analyses, particularly with methods like *Relate* and haplotype-based tests (*iHS*, *Rsb*, and *XP-EHH*). For the genome scan using *PBS*, which is more robust with small samples (*90*), the selection signals may not have been detected due to the similarity of *P. simium* sequences to those of Brazilian and Colombian *P. vivax* (*21*), the reference populations used in the test. Therefore, additional *P. simium* samples are necessary to identify such signals.

We found no evidence of selection in *P. simium* genes associated with resistance to antimalarial treatments, primarily chloroquine and primaquine. For *P. vivax*, no chloroquine resistance was reported in Latin America before 2000, and its spread across the continent remains unclear (*91*, *92*). Additionally, no *in vivo* resistance to primaquine has been reported (*92*). Consequently, it is possible that *P. simium* is not resistant to these treatments, which would explain the absence of selection evidence for resistance-related genes. Similarly, Brasil *et al.* (*5*) reported no treatment resistance in humans infected with *P. simium*. However, the limited sample size (n=19) may have been insufficient to detect such signals.

Noteworthy, we detected significant positive selection signals in 73 genes, including *PvIMC1h*, *PvG377*, and *PvGRASP* (Table S3). These genes may have played a role in *P. simium* adaptation to the mosquito and/or primate host, but their function is currently unknown. Functional analyses could provide additional insights into *P. simium* adaptive processes.

In conclusion, zoonotic malaria is a significant issue in Latin America, as highlighted by a recent *P. simium* outbreak in the Atlantic Forest of Rio de Janeiro, Brazil. This underscores the potential role of NHPs malaria reservoirs and emphasizes the need of extensive research on their genetic diversity, evolutionary history, and zoonotic potential. Furthermore, the health and conservation impacts of these infections on NHPs, particularly endangered species in the Atlantic Forest, remain largely unexplored. Our results, based on 646 NHP samples from five Latin American countries, provide the first evidence of natural *P. vivax* infections in Colombian NHPs, highlighting the possibility of ongoing host switches and their relevance for malaria elimination strategies. The genetic analyses revealed recent admixture events and gene flow between *P. simium* and *P. vivax* populations, emphasizing the porous boundaries between these species and their adaptive processes. Using various population genomic approaches, we consistently demonstrated that *P. simium* forms a separate cluster more closely related to American *P. vivax* populations. We then identified two genetic clusters within *P. simium*: one predominantly in Rio de Janeiro state and another in neighbouring São Paulo state. The close genetic relationship between *P. simium* and human *P. vivax* suggests recent host transfers, with divergence occurring in the last 30 years and migration events dating back to 100-200 years, likely involving a host jump from Mexican *P. vivax* to Brazilian *P. simium*. Despite the limited sample size, which may have hindered the detection of some selection signals, we identified several genes potentially involved in *P. simium* adaptation to NHPs, human red blood cells, and mosquito vectors. Our findings question the distinction between *P. simium* and *P. vivax* as separate species because both parasites can infect humans and Neotropical NHPs and produce hybrids, and suggest that *P. simium* may represent distinct *P. vivax* populations. The results presented in this study have profound implications for malaria eradication efforts in Latin America. By highlighting the potential NHPs role as zoonotic malaria reservoirs and the porous genetic boundaries between *P. vivax* and *P. simium*, our study underscores the need to incorporate zoonotic dynamics into malaria control strategies, while promoting coexistence between humans and NHPs.

## Materials and methods

### *P. simium* sample collection and ethical statements

In French Guiana, non-human vertebrate samples were collected and used in accordance with an international CITES permit (Convention on International Trade in Endangered Species of Wild Fauna and Flora; permit FR973A) according to the French legislation. Following the sharing policy in French Guiana, non-human mammal samples are registered in the collection JAGUARS (https://kwata.net/gestion-collection-biologique/, CITES reference: FR973A) supported by the Kwata NGO (accredited by the French Ministry of the Environment and the Prefecture of French Guiana, Agreement R03-2019-06-19-13), Institut Pasteur de la Guyane, DGTM Guyane, Collectivité Territoriale de la Guyane, and validated by the French Guianan prefectoral decree n°2012/110.

In Costa Rica, the animal study was reviewed and approved by the Institutional Committee for the Care and Use of Animals (Comité Institucional para el Cuidado y Uso de los Animales) of the Universidad de Costa Rica, and adhered to the Costa Rica legal requirements (collection permit number: MINAET-SINAC, Costa Rica: 042-2012-SINAC).

In Brazil, the collected animals were dead and part of the epidemiology study of the Rio Grande do Sul State Health Secretaria, with a license issued by the Ministry of Health.

In Colombia, ethical approvals for the collection of fecal and blood samples as well as the collection of vectors were obtained by Universidad de los Andes, the National Environmental Licensing Authority of Colombia (ANLA) and the Centers for Disease Control and Prevention (permits numbers: 2017025578-1-000, 2017043863-1-000, 2017065795-1-000, 2017013727-1-000, 2017052943-1-000, 2017081458-1-000, 2017108650-1-000).

In Argentina, permits were obtained from National Parks to conduct this research (IF 2022 85832064 APN DRNEA # APNAC PROYECT NUMBER 541 and IF 2022 85832064 APN DRNEA # APNAC PROYECT NUMBER 544).

### DNA extraction and *Plasmodium* screening

For each of the 646 samples collected from NHPs in Latin America, genomic DNA was extracted using the Qiagen DNeasy Blood and Tissue Kit according to the manufacturer’s recommendations. *P. simium* and *P. vivax* samples were identified by amplification of *Plasmodium cytochrome b* using nested PCR, as described in Prugnolle *et al.* (*31*). The reaction products were visualized on 1.5% agarose gels stained with EZ-vision.

For *P. simium* or *P. vivax* positive samples, selective whole-genome amplification (sWGA) was performed to enrich submicroscopic levels of *Plasmodium* DNA, following the protocol described by Cowell *et al.* (*93*). This technique preferentially amplifies *P. vivax* and *P. simium* genomes from a set of target DNAs and reduces host DNA contamination. For each sample, DNA amplification was carried out using the strand-displacing phi29 DNA polymerase and *P. vivax*-specific primers that target short (6 to 12 nucleotides) motifs commonly found in the *P. vivax* genome (PvSet1 (*93*, *94*)). Although *P. vivax*-specific, these primers have already been used successfully to amplify other *Plasmodium* species phylogenetically close to *P. vivax* such as *P. vivax-like* identified in African great apes(*36*). About 30 ng of input DNA was added to a 50 µL reaction mixture containing 3.5 μM of each sWGA primer, 30 U of phi29 DNA polymerase enzyme (New England Biolabs), 1× phi29 buffer (New England Biolabs), 4 mM deoxynucleotide triphosphates (Invitrogen), 1% bovine serum albumin, and sterile water. DNA was amplified in a thermal cycler with the following program: a ramp down from 35° to 30°C (10 min per degree), 16 hours at 30°C, 10 min at 65°C, and hold at 4°C. For each sample, the products of the two amplifications (one per primer set) were purified with AMPure XP beads (Beckman Coulter) at a 1:1 ratio according to the manufacturer’s recommendations and pooled at equimolar concentrations. Samples with the highest concentration of parasite genome were selected after qPCR analysis using a Roche LightCycler 96 with the following program: 95°C for 10 minutes; 40 cycles of 95°C for 15 seconds, 60°C for 20 seconds, and 72°C for 20 seconds; 95°C for 10 seconds; and 55°C for 1 minute. Each sWGA library was prepared using the two pooled amplification products and the Nextera XT DNA kit (Illumina), following the manufacturer’s protocol. Then, samples were pooled, clustered, and sequenced on one lane of a Illumina Novaseq-6000 S4 with PE150 with 2 × 150-bp paired-end reads.

### Short-read mapping, SNP calling, and data compilation

Three newly sequenced *P. vivax* isolates collected in Colombian monkeys were added to a compilation of previously published genomic datasets: *i*) 621 *P. vivax* samples from Daron *et al.* (*36*), Benavente *et al.* (*87*), the MalariaGEN *P. vivax* Genome Variation project (*95*), Mourier *et al.* (*20*), and Lefebvre *et al.* (*37*); *ii*) 31 *P. simium* samples from Mourier *et al.* (*20*), de Oliveira *et al.* (*21*), and Ibrahim *et al.* (*19*); and *iii*) two *P. vivax-like* genomes collected on Nigeria-Cameroon chimpanzees (*Pan troglodytes ellioti*), from Loy *et al.* (*96*) (Fig. S1). Short read archives (SRA) from the literature were retrieved from NCBI (accession numbers in Table S1).

Short reads were trimmed to remove potential lingering adapters and preprocessed to eliminate low-quality reads (*−quality-cutoff = 30*) using the *cutadapt* program (*97*). Reads shorter than 50 bp and containing “N” were discarded (*−minimumlength = 50 –max-n = 0*). Cleaned paired-end reads were mapped to the *P. vivax* reference genome PVP01 (*98*) using *bwa-mem* (*99*). Duplicate reads were marked using the *MarkDuplicates* tool from the *Picard tools* v2.5.0 (broadinstitute.github.io/picard/) with default options. Local realignment around indels was performed using the *IndelRealigner* tool from *Genome Analysis Toolkit* (*GATK* (*100*), v3.8.0). Variants were called using the *HaplotypeCaller* module in *GATK* with default options. Lastly, the different isolated variant call format (VCF) files were merged using the *GATK* module *CombineGVCFs*.

### Data filtering

For all samples (*P. vivax* and *P. simium*) from Mourier *et al.* (*20*), de Oliveira *et al.* (*21*), and Ibrahim *et al.* (*19*), whether *P. vivax* or *P. simium*, the same filters used in Lefebvre *et al.* (*37*) were applied. Thus samples with >50% missing data were removed. As several strains can infect the same host, the within-host infection complexity was assessed with the *F_WS_* metric (*101*), calculated using *vcfdo* (github.com/IDEELResearch/vcfdo; last accessed July 2022). Samples with mono-clonal infections (*i.e.* F_WS_ >0.95) were kept (Table S4). Highly related samples and clones could generate spurious signals of population structure, biased estimators of population genetic variation, and violated the assumptions of the model-based population genetic approaches used in this study (*102*). Therefore, the relatedness between haploid genotype pairs was measured by estimating the pairwise fraction of the genome in IBD using the *hmmIBD* program (*42*), with the default parameters for recombination and genotyping error rates, and using the allele frequencies estimated by the program. Within each country and species, isolate pairs that shared >50% of IBD were considered highly related. Only the strain with the lowest amount of missing data was retained in each family of related samples.

The final dataset included 429 samples: 404 *P. vivax* isolates, 19 *P. simium* isolates, 4 *P. vivax* isolates from NHPs, and 2 *P. vivax-like* isolates used as outgroup species in phylogenetic analyses.

For the VCF with 426 samples (without the three newly sequenced samples), only bi-allelic SNPs with less than 50% missing data and a genotype quality (GQ) score above 30 were kept. SNPs with a minimum average depth coverage of 5x and a maximum average depth of 152x on all samples were kept. Singletons were removed because they can be sequencing errors. This dataset based on hard filters (*dataset_HF*) included 819,652 SNPs with a mean density of 38.29 SNPs per kilobase (Fig. S1).

For the *ANGSD* software suite that uses the genotype likelihood (GL or GQ) rather than the actual genotype calling, all sites with a genotype quality >20, a maximum depth of 155x per sample, a minimum total depth of 5x on all samples, and present in at least 5 individuals were kept. Only bi-allelic SNPs with a *p-value* of 1e-6 were kept. A total of 20,844,131 sites from the core genome were considered, including 1,286,079 variants (65.22 SNPs/kb). This approach allowed considering low to very low coverage samples (*33*, *103*) and led to a dataset that included all 429 samples (*dataset _GL*). All filtration steps for both datasets are detailed in Fig. S1.

### Population structure and relationships among populations

The principal component analysis (PCA) and the individual genetic ancestry analyses were carried out with *ANGSD* v.040 and *PCAngsd* v0.98, after selecting only biallelic SNPs present in the *P. vivax* core genome and excluding SNPs with a minor allele frequency (MAF) <5% from *dataset_GL*. Variants were LD-pruned to obtain a set of unlinked variants using *ngsld* v1.1.1(*104*), with a threshold of *r*²=0.5 and windows of 0.5 kb. In total, these analyses included 247,890 SNPs for 425 individuals (the *P. vivax-like* outgroup was not considered). For the ancestry analyses, using *PCAngsd*, a number of clusters (*K*) ranging from 2 to 8 were tested. The best K value was inferred based on the broken-stick eigenvalues plus one (Fig. S2), following the recommendations of Meisner and Albrechtsen (*34*). The results were post-processed using *pong* v1.5 (*105*) to compute the ancestry proportions.

The maximum likelihood (ML) phylogenetic tree was obtained with *IQ-TREE* (*35*) using the best-fitted model determined by *ModelFinder* (*106*). Considering the *dataset_HF,* only SNPs with ≤10% missing data and samples with <50% missing data were retained, resulting in 48,070 SNPs in the core genome for 404 samples. As the dataset was composed only of SNPs and no invariant sites, the ascertainment bias correction was added to the tested models, following the *IQ-TREE* user guide recommendations. According to the Akaike information criterion (*107*), the best model of nucleotide evolution was the general time reversible (GTR) model that integrates unequal rates and unequal base frequency (*108*). Node support was estimated using the Ultrafast Bootstrap Approximation (UFboot) (*40*) and SH-aLRT methods (*39*). Nodes were considered as highly supported only if SH-aLRT ≥ 80% and UFboot ≥ 95%, following the thresholds by Guindon *et al.* (*39*) and Minh *et al.* (*40*).

PCA and genetic ancestry analysis were performed focusing only on the *P. simium* samples to investigate the genetic substructure within this group. As these analyses require a data set with unlinked variants, SNPs from the *dataset_GL* (with only *P. simium* samples) were LD-pruned with *ngsld* v1.1.1 (*104*), with a threshold of *r*²=0.5 and windows of 0.5 kb. In total, these analyses included 44,911 SNPs for 19 individuals. The PCA was carried out with *ANGSD* v.040 and *PCAngsd* v0.98, after selecting only biallelic SNPs present in the *P. vivax* core genome and excluding singletons. Then, individual genetic ancestry was estimated using the *PCAngsd*. Different cluster (*K*) numbers were tested from 1 to 5. The optimal *K* value was estimated based on the broken-stick eigenvalues plus one (Fig. S2), following the recommendations of Meisner and Albrechtsenc(*34*), and the plots were generated with *pong* (*105*).

For the pairwise-IBD network, the complete *dataset_HF*, composed of 819,652 SNPs and 426 samples, was used. Pairwise-IBD values among pairs of isolates were estimated using the *hmmIBD* program (*109*), with the default parameters for recombination and genotyping error rates, and using the allele frequencies estimated by the program. The pairwise-IBD relationships among isolates were visualized as a network that displayed only pairwise IBD relationships with values ≥ 5%. The network was plotted using the *igraph* package in R (*41*) and default options.

Population graphs were built using *TreeMix* (*44*) and *AdmixtureBayes* (*43*). First, the *dataset_HF* was pruned using PLINKc(*110*) with a threshold of *r*²=0.5, a sliding window of 50 SNPs and a step of 10 SNPs (396,835 SNPs remaining). All SNPs with missing data were removed. The final dataset used in these analyses included 11,907 SNPs for 18 ingroup populations and one outgroup population (*P. vivax-like*).

The number of migration events (*m*) in *TreeMix* that best fitted the data was calculated by running the analysis 15 times for each *m* value, with *m* ranging from 1 to 10. The optimal *m* value was estimated using the *OptM* R package (*111*) and the Evanno method (Fig. S5). Then, a consensus tree with bootstrap node support was obtained by running 100 replicates of *TreeMix* and postprocessing them using the *BITE* R-package (*112*) for the *m* optimal values, following Daron *et al.* (*36*) and Lefebvre *et al.* (*113*).

For the *AdmixtureBayes* analysis, three independent runs were performed, each one including 40 coupled Monte Carlo Markov Chains (MCMC) and 250,000 steps. The results of these chains were analyzed after a burn-in of 50% and a thinning step of 10, using the default parameters. Convergence across the trees was assessed using the Gelman-Rubin (GR) statistics for the models parameters (i.e., posterior probability, branch lengths and number of admixtures), as recommended by Nielsen *et al.* (*43*). Runs were deemed convergent if the GR statistics for the model parameters converged toward the value of 1 along the MCMC steps (Fig. S4). The retained network was common among the three best networks in each chain that converged, based on their posterior probabilities.

The admixture *f_4_*-statistics were computed with *ADMIXTOOLS 2* v2.0.0(*46*) and the same dataset used for *TreeMix* and *AdmixtureBayes*,.

### Demographic History of American *P. simium*

The nucleotide diversity (π) values were estimated using *pixy* v1.2.7 (*114*), in a non-overlapping sliding window of 500 bp along the genome based on the *dataset_GL*. The sample size was standardized (*i.e.*, 10 randomly chosen isolates for each population) to obtain values that could be compared.

*Relate* (*47*) was used to infer historical changes in *N_e_* and to estimate the divergence time between populations. The ancestral/derived states of 1,267,897 sites from the *dataset_GL* were defined using the consensus sequence of the outgroup (*P. vivax-like*). Only the core genome regions as defined in Daron *et al.* (*36*) were considered, the rest was masked. As the *P. simium* samples were not enough (n=19) to generate a *P. simium*-specific genetic map, the *P. vivax* genetic map from Lefebvre *et al.* (*37*) was used with a mutation rate of 6.43×10^-9^ mutations/site/generation and a generation time of 5.5 generations/years(*36*). Coalescence rates were calculated from 10 to 1,000,000 generations ago (options *--bins 0,7,0.25*).

Then, MSMC-IM (*48*), based on the coalescent rates estimated within and between two populations from *Relate* (*47*), was used to fit an isolation-with-migration (IM) model. The scripts to convert *Relate* results to *MSMC-IM* were kindly provided by E. Patin and D. Liu from the Human Evolutionary Genetics team at the Institut Pasteur, Paris, France. The migration rates were inferred between *P. simium* clusters and two *P. vivax* populations (Brazil and Mexico). These *P. vivax* populations were selected based on their known evolutionary history and our population structure results. Brazilian and Mexican *P. vivax* were included due to their geographic proximity to *P. simium*, and genetic similarity to *P. simium*, respectively. The mutation rate was the one used for *Relate*, and the population sizes were derived from the *Relate* results.

### Positive selection detection

Given the low sample size in the two *P. simium* genetic clusters (n = 13 and n = 6), they were combined for the selection scan analyses with *rehh* (*115*) and *Relate* (*47*), following Hupalo *et al.* (*38*) and Lefebvre *et al.* (*113*).

The haplotype-based tests (*iHS, XP-EHH,* and *Rsb*) were used to detect evidence of recent selective sweeps, using *rehh* (*115*) and *dataset_HF*. The significance threshold was set at -log(*p-value*) = 4, as recommended by Gautier *et al.*(*115*). For *XP-EHH* and *Rsb*, the *P. simium* samples were compared with Brazilian *P. vivax* samples and only SNPs with negative standardized values (*i.e.*, indicating positive selection for the *P. simium* cluster rather than for *P. vivax*) were considered. The ancestral/derived states of 378,441 SNPs from *dataset_HF* were defined using the consensus sequence of the two outgroup *P. vivax-like* isolates. The Brazilian population was chosen due to its larger sample size (n=29). The Mexican population did not have a large enough sample size (n=9) to be used as a reference population to detect selection signals (*116*).

The *Relate* program used to estimate the coalescent rates in the demographic section was also used to detect positive selection along the genome by identifying genomic regions with significantly higher coalescent rate values than the rest of the genome. The significance threshold was set at -log(*p-value*) = 4, to take into account the multiple tests.

The population branch-site (*PBS*) values were calculated for each *P. simium* cluster, with *ANGSD* v0.40 in sliding windows of the genome (windows of 1 kb with a sliding step of 500 pb), based on *dataset_GL*. The outgroup populations were *P. vivax* from humans in Brazil and Colombia. All windows with <20 SNPs were removed to avoid extreme values caused by a low SNP number. The 0.1% most extreme values were considered evidence that the genomic regions displayed signs of selection in our *P. simium* clusters.

Once selection signals were detected, the identified genes were annotated using the general feature format (GFF) file (*PlasmoDB-50_PvivaxP01.gff*, available on *PlasmoDB*) and the intersect function of *BEDtools* v 2.26.0 (*117*). Additional information (*e.g.*, gene name, function, and biological process) was retrieved from *PlasmoDB* (plasmodb.org, accessed in April 2024).

## Acknowledgments

The authors acknowledge the ISO 9001-certified IRD i-Trop HPC (member of the South Green Platform) at IRD Montpellier for providing HPC resources that contributed to the research results reported in this paper. URL: https://bioinfo.ird.fr/- http://www.southgreen.fr. The bioinformatics analyses were also performed on the Core Cluster of the Institut Français de Bioinformatique (IFB) (ANR-11-INBS-0013). We thank Olivier Duron for facilitating the connection with Benoit De Thoisy, who provided the samples from French Guiana. We would like to thank Josquin Daron for providing the scripts for sample mapping and calling. We also thank Leo Speidel for his guidance on *Relate,* and Dang Liu and Etienne Patin for providing the scripts to convert results from *Relate* to *MSMC-IM*. Finally, we thank Elisabetta Andermarcher for proofreading the manuscript and supplementary material.

## Funding

project ORA in the LabEx CEBA, ANR-10-LABX-25-01 (VR)

JCJC GENAD, ANR-20-CE35-0003 (VR)

MICETRAL, ANR-19-CE350010 (FP)

the United States (U.S.) Agency for International Development and the U.S. Centers for Disease Control and Prevention (CDC) for the project entitled "Evaluation of ZIKV potential to establish a sylvatic transmission cycle in Colombia" (CGR, AL, SR)

## Author contributions

Conception: V.R., F.P., M.C.F. and M.J.M.L.

Funding acquisition: V.R., M.C.F., F.P.

Biological data acquisition and management: V.R., B.DT., C.G., S.R., A.L., A.C., J.A.B., A.S.C., E.dS.,R.H., C.M.DdS., E.V.

Sequence data acquisition: F.D. and C.A.

Method development and data analysis: M.J.M.L., V.R., F.P., and M.C.F.

Interpretation of the results: V.R., F.P., M.C.F., and M.J.M.L.

Drafting of the manuscript: M.J.M.L., V.R., M.C.F., and F.P.

Reviewing and editing of the manuscript: V.R., F.P., M.C.F., and M.J.M.L. with the contribution from all the co-authors.

## Competing interests

The authors declare that they have no competing interests.

## Data availability

The sequences of the newly sequenced and analyzed simian *P. vivax* samples have been deposited on NCBI (Bioproject PRJNA116416). SNP data files (VCF format) produced in this study as well as the related metadata and documentation are available in DataSuds repository (IRD, France) at https://doi.org/10.23708/RCOGR6.

The scripts used in this study are available in this github repository: https://github.com/MargauxLefebvre/EvoHistory_Simium

## Notes

### Competing Interest Statement

The authors have declared no competing interest.

### Summary of Updates

In this revised version, we better account for the variability in P. simium data coverage in the discussion and by using genotype likelihood-based tools, such as ANGSD and PCAngsd, for population structure analysis.

