## Supplementary material for "Genomic insights into host shifts between *Plasmodium vivax* and *Plasmodium simium* in Latin America": Fig. S

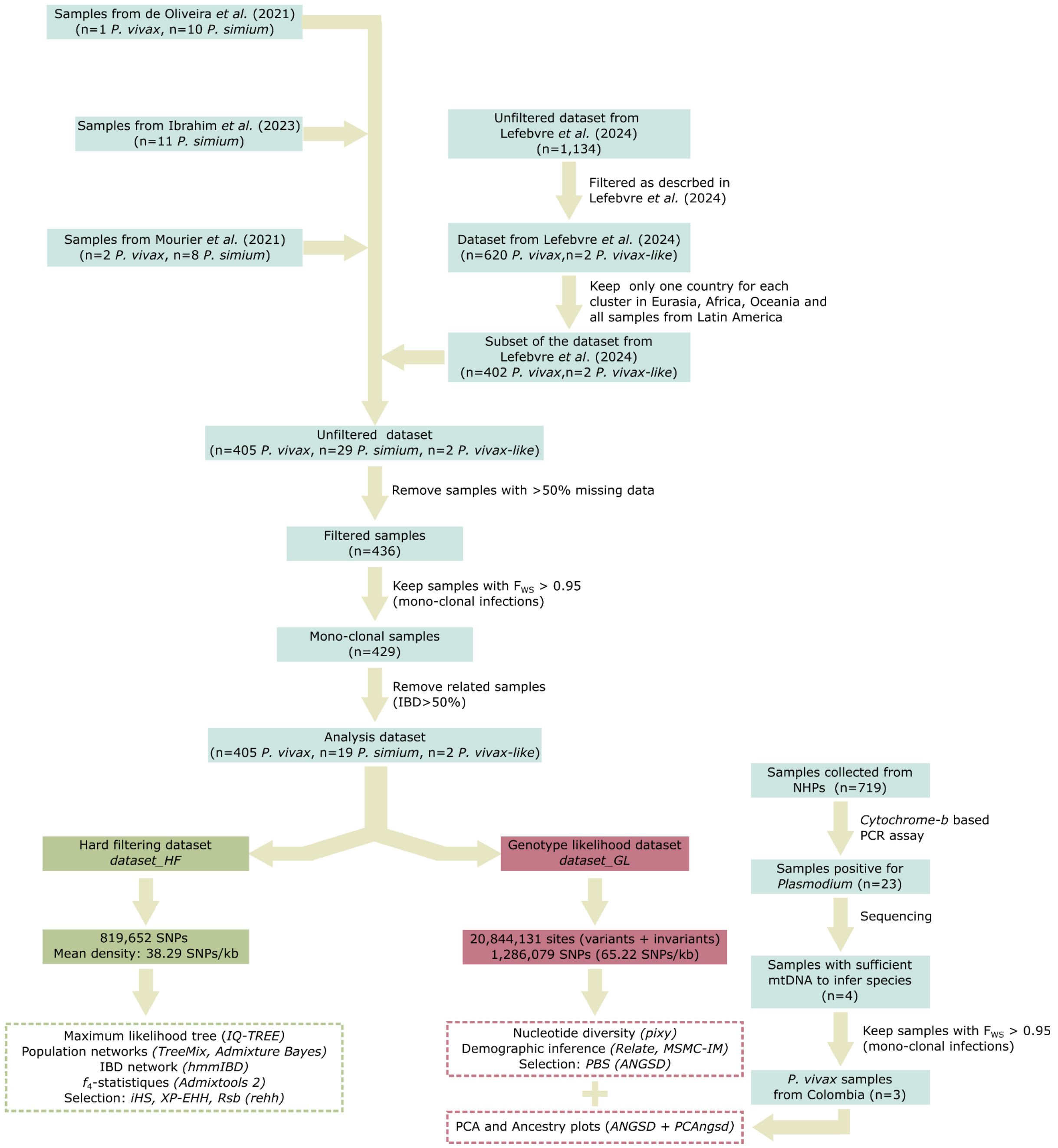


**Fig. S1.**

**Filtering and dataset processing.** Each box in the diagram represents a specific filtering step and indicates the number of remaining samples or SNPs. Key filtering options are also specified. The final steps (dotted line boxes) show the analyses performed using the different datasets.

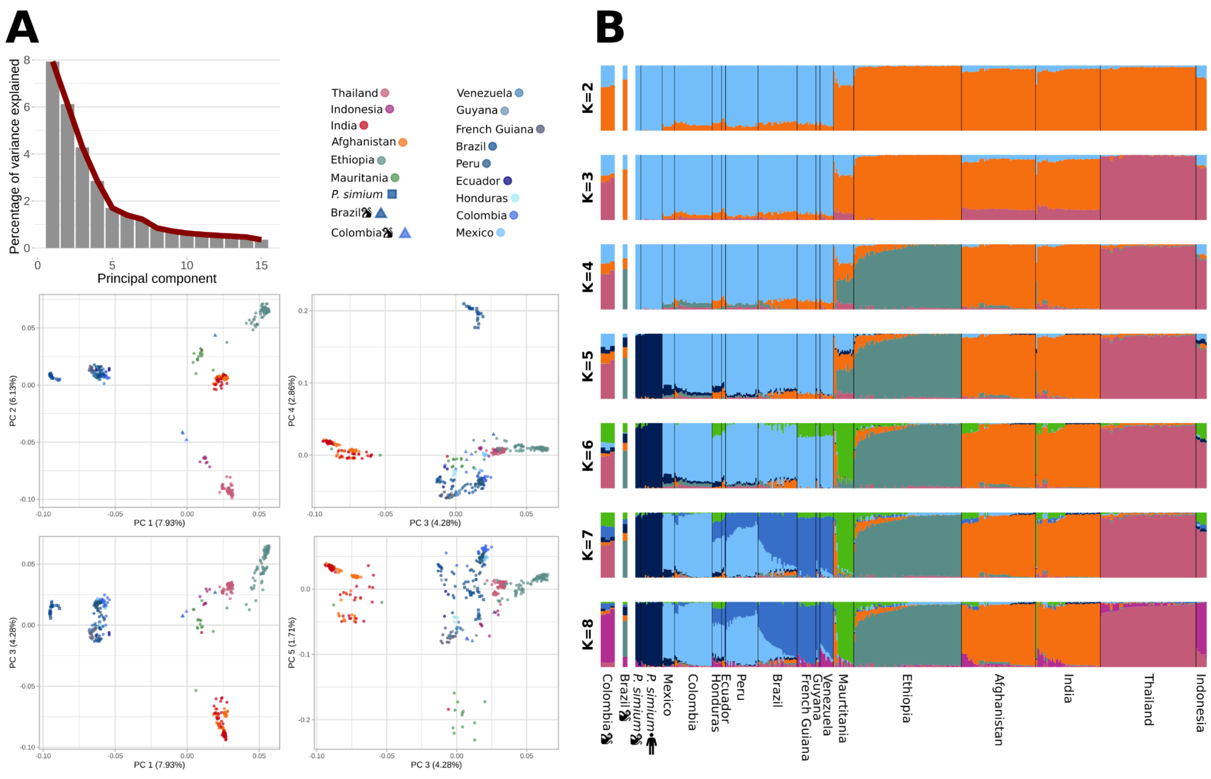


**Fig. S2.**

**(A)** Principal component analysis (PCA) of 408 *P. vivax* and 19 *P. simium* strains based on the genotype likelihood of 247,890 unlinked SNPs. The bar plot (upper panel) shows the percentage of variance explained by the first 15 principal components (PC). The optimal PC number is 5, according to the elbow (broken-stick) method. PCA plots for PC 1 to 5 (lower panels). **(B)** Genetic ancestry of *P. vivax* and *P. simium* samples worldwide estimated with *PCAngsd* (K=2 to K=8). The number (K) of clusters tested is specified on the left. According to Meisner and Albrechtsen (*34*), the best K is determined by 1+ D, where D is the optimal number of principal components. As panel (A) shows that the best D would be 5, the best K value would be 6. The monkey pictogram indicates *P. vivax* and *P. simium* isolates sampled in monkeys from Latin America. For *P. simium* samples, the human pictogram indicates samples from humans.

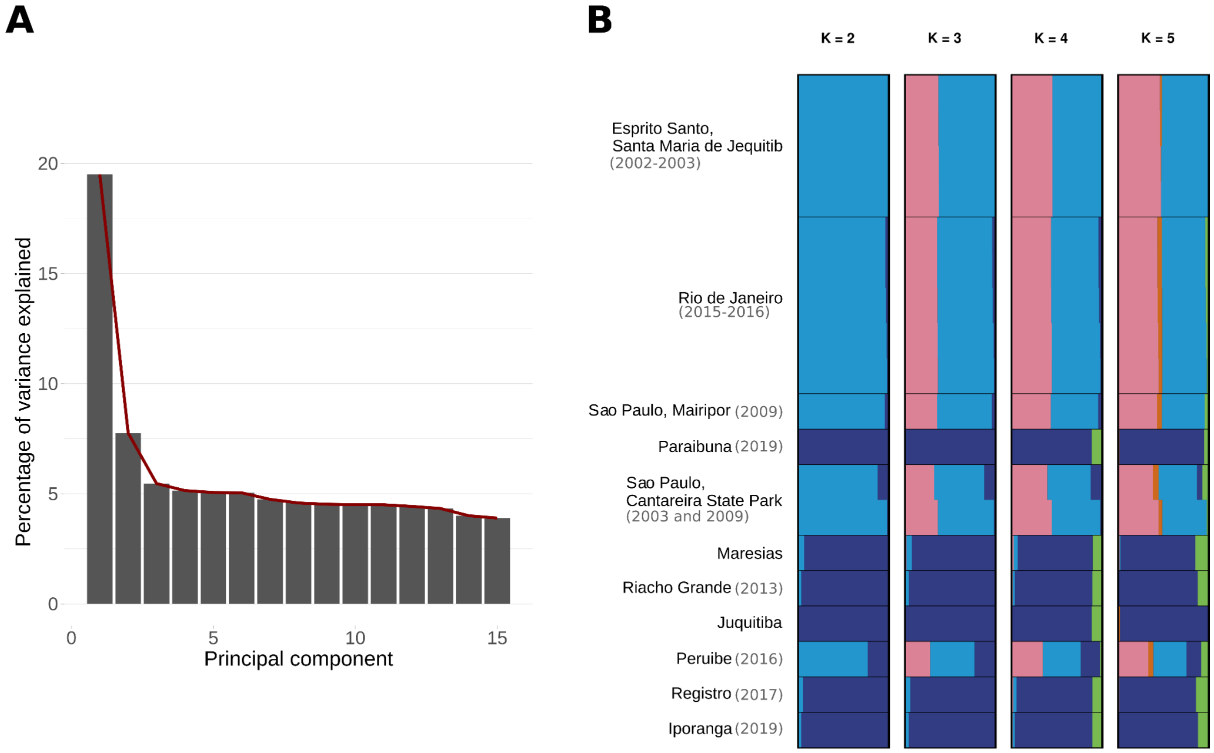


**Fig. S3.**

**(A)** Principal component analysis (PCA) of 19 *P. simium* strains based on the genotype likelihood of 44,911 unlinked SNPs. The bar plot shows the percentage of variance explained by the first 15 principal components (PC). The optimal PC number is 1, according to the elbow (broken-stick) method. **(B)** Genetic ancestry of *P. simium* samples worldwide estimated with *PCAngsd* (K=2 to K=8). The number (K) of clusters tested is specified on the top. According to Meisner and Albrechtsen (*34*), the best K is determined by 1+ D, where D is the optimal number of principal components. As panel (A) shows that the best D would be 1, the best K value would be 2.

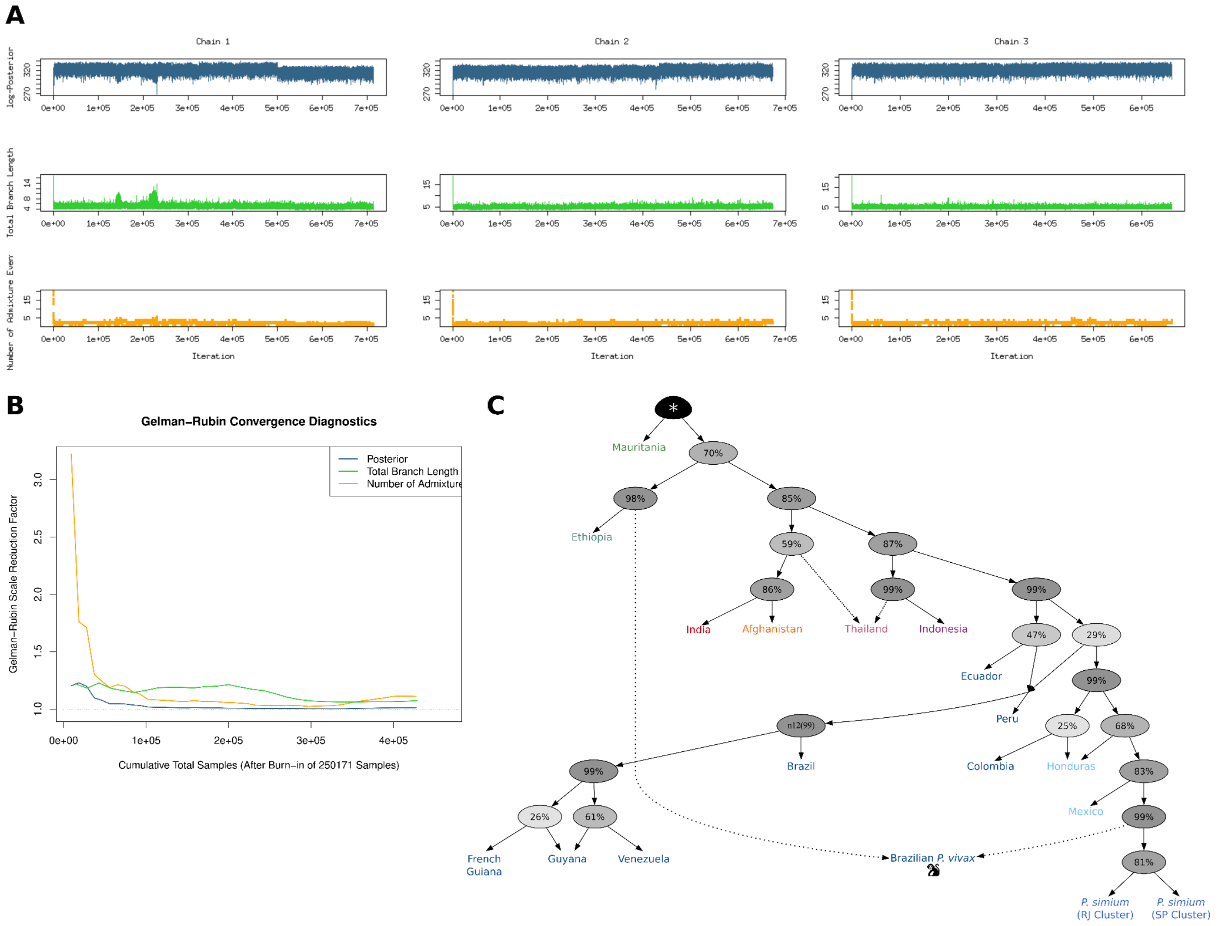


**Fig. S4.**

**Identification of the optimal number of admixture events for *P. vivax* and *P. simium* populations and consensus topology computed with *AdmixtureBayes*. (A)** Trace plots of the posterior probability, total branch length, and number of admixture events for each chain. **(B)** Plot of the Gelman-Rubin convergence diagnostics on the chains for the indicated summary statistics after a burn-in fraction of 50%. The trends for the different statistics approaching 1 indicate convergences of the MCMC chains. **(C)** Consensus population graph generated by combining nodes with a posterior probability >25%. Dotted lines represent admixture events. The percentages at the nodes indicate the posterior probability that the true graph has a node with the same descendants. The monkey pictogram indicates the *P. vivax* isolate sampled in a monkey from Latin America.

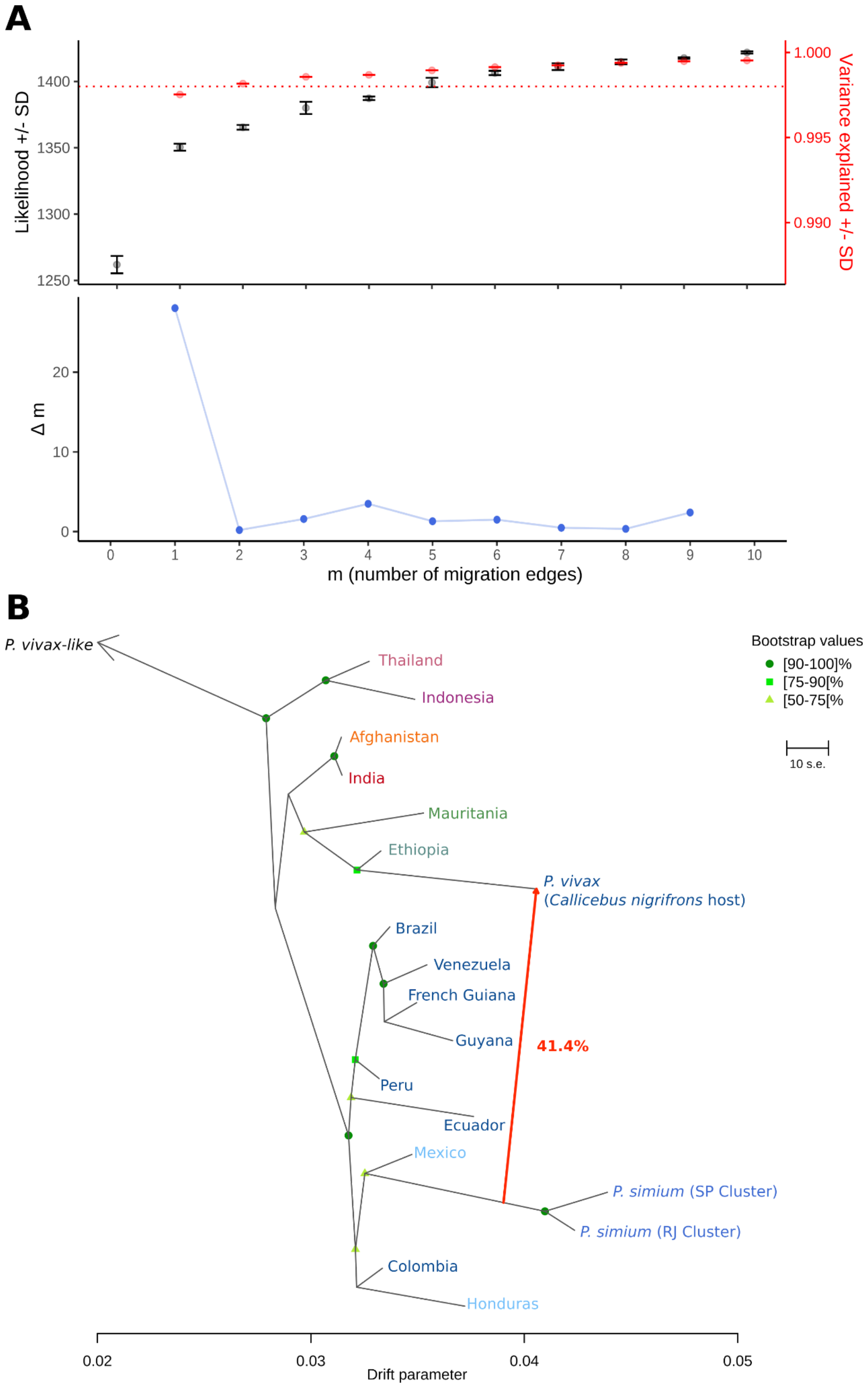


**Fig. S5.**

**Identification of the optimal number of migration edges (*m*) for 16 *P. vivax* and 2 *P. simium* populations and the population consensus tree topology for m=1 estimated with *TreeMix*. (A)** Changes in the mean likelihood score (± SD) and the average total fraction of the genetic variance explained (± SD) in function of the number of migration edges considered in the *TreeMix* analysis (upper panel). Second-order rate of change in likelihood (*Δm*) across migration edges (*m*) values, from 0 to 10 (lower panel). The inflection point was observed at m=1. **(B)** *TreeMix* tree of *P. vivax* and *P. simium* populations with one migration edge (arrow), rooted with *P. vivax-like*. The scale bar shows ten times the mean standard error (s.e.). The migration weight is indicated next to the arrow.

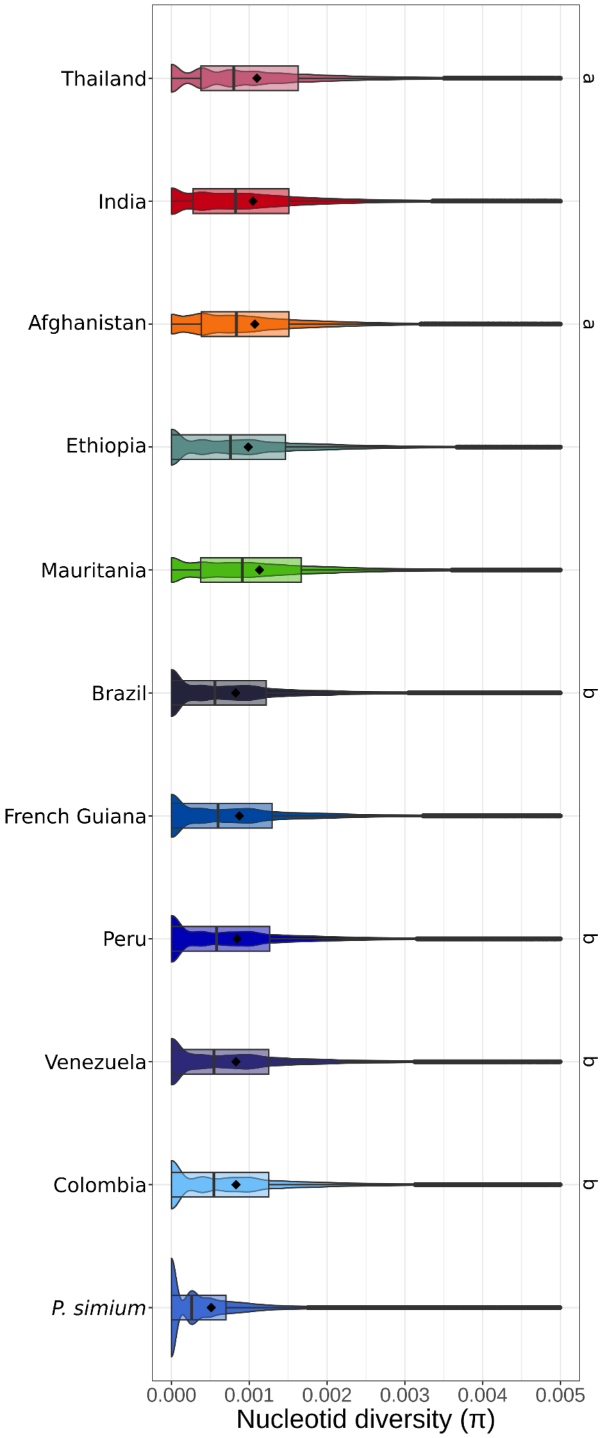


**Fig. S6.**

**Genetic diversity of *P. vivax* and *P. simium* populations.** Genome-wide distributions of the nucleotide diversity (π) values in 500 bp non-overlapping windows along the core genome for populations with >10 samples. The boxplot shows the three quartiles (the first at 25%, the median at 50%, and the third at 75%), and the diamond indicates the mean. The same letter (on the right) indicates no significant difference between the distributions (Wilcoxon test and Bonferroni correction). If there is no letter, the distribution significantly differs from all the other distributions (*p-value* < 0.001). Note that *P. simium* stands out from other *P. vivax* populations by having a considerably lower nucleotide diversity.

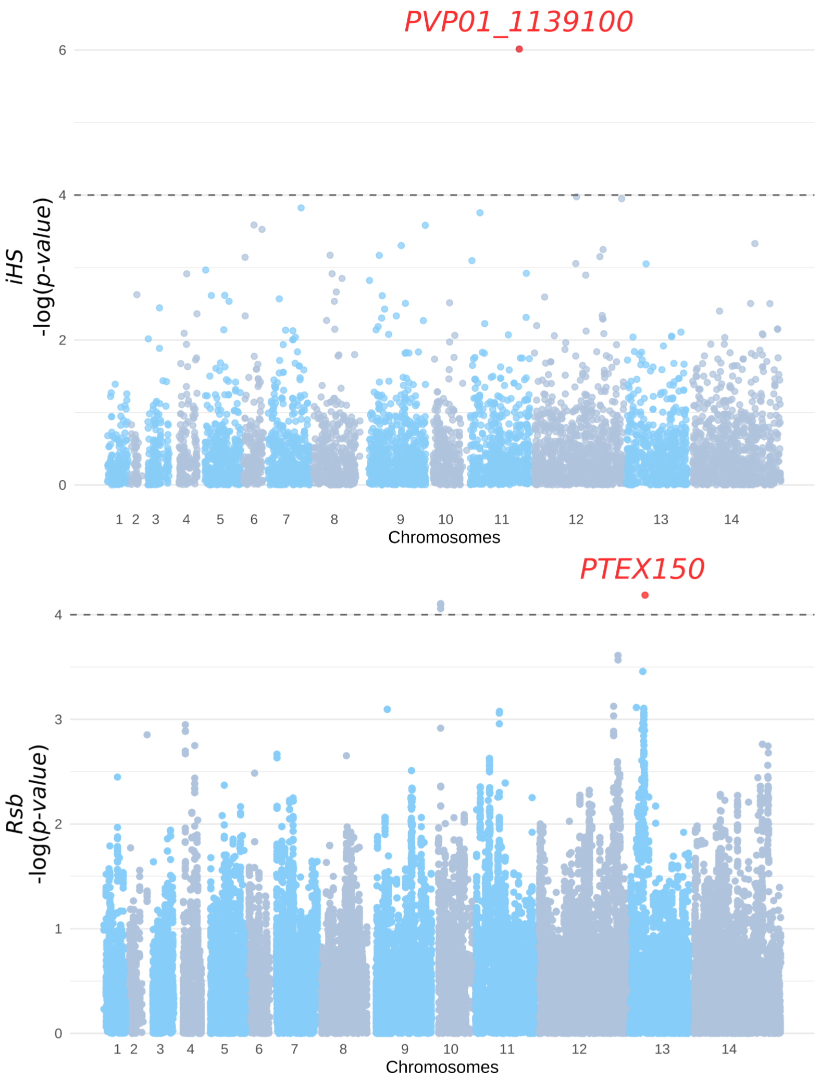


**Fig. S7.**

**Haplotype-based test of positive selection relying on the *iHS* (top) and *Rsb* (bottom) statistics.** Dotted lines represent the significance threshold value -log(*p-value*) = 4. The red points are SNPs detected as significantly affected by a selective sweep in *P. simium* (negative values for *Rsb*).

**Table S1.**

**Sample metadata.** The NCBI SSR-ID, bioproject, biosample, and source are indicated for each sample. When available, the latitude and longitude are specified. NA, information not available. The QC column indicates whether samples successfully passed the quality control (QC) and were included in the final dataset for analyses. For samples that did not pass the QC, the reasons are listed in the "Reasons_QC" column as follows: "Missing_data" for samples with >50% missing data, "Low_Fws" for samples removed due to multi-clonal infections, and "High_IBD" for samples that are related to those kept in the analysis dataset. For additional details, please refer to the Materials and Methods section and Fig. S1.

**Table S2.**

**Samples from American monkeys screened for *Plasmodium* infection.** For each sample, the host species, sampled organ(s), country, region, and collection date are provided. "NA" indicates data not available. The "PCR cytB" and "PCR Liu" columns report the results of nested PCR targeting *Plasmodium* cytochrome *b* and Liu gene regions, respectively. The "Sequenced" column indicates whether samples were submitted for whole-genome sequencing.

**Table S3.**

**List of genes in which evidence of positive selection was observed in *P. simium* using *iHS*, *XP-EHH*, *Rsb*, *PBS* and *Relate*.** The “selection_test” column indicates the test with which this gene was detected. For PBS, it is also specified in which cluster: “PBS_RJ” for the RJ cluster and “PBS_SP” for the SP cluster. Genes in bold are discussed in this study.

**Table S4.**

**$F_{WS}$ values for samples added to the dataset from Lefebvre *et al.* (*37*).** NA, information not available for samples with not enough data to infer the $F_{WS}$ values.
